# FIGL1 attenuates meiotic interhomolog repair and is counteracted by RAD51 paralog XRCC2 and chromosome axis protein ASY1 during meiosis

**DOI:** 10.1101/2024.04.25.591231

**Authors:** Côme Emmenecker, Simine Pakzad, Fatou Ture, Julie Guerin, Aurelie Hurel, Aurelie Chambon, Chloe Girard, Raphael Mercier, Rajeev Kumar

## Abstract

Two recombinases, RAD51 and DMC1, catalyze meiotic break repair to ensure crossovers (COs) between homologous chromosomes (interhomolog) rather than between sisters (intersister). FIDGETIN-LIKE-1 (FIGL1) downregulates both recombinases. However, the understanding of FIGL1 functions in meiotic repair remains limited. Here, we discover new genetic interactions of *Arabidopsis thaliana FIGL1* that are important *in vivo* determinants of meiotic repair outcome. In *figl1*, compromising the RAD51-dependent repair by either losing RAD51 paralogs (RAD51B or XRCC2) or RAD54 or inhibiting RAD51’s catalytic activity results in either unrepaired breaks or meiotic CO defects. Further, XRCC2 physically interacts with FIGL1 and partially counteracts FIGL1 for RAD51 focus formation. Our data support that RAD51-mediated repair mechanisms compensate for the FIGL1 dysfunction. FIGL1 is dispensable for intersister repair in *dmc1* but is essential for meiotic repair completion in mutants with impaired DMC1 functions and interhomolog bias such as *asy1*. We show that FIGL1 attenuates interhomolog repair, and ASY1 counteracts FIGL1 to promote interhomolog recombination.

## Introduction

During meiosis, the repair of DNA double-stranded breaks (DSBs) by homologous recombination (HR) yields crossovers (COs) and non-crossovers (NCOs) (1, 2). Meiotic COs between homologous chromosomes (interhomolog) rather than between sister chromatids (intersister) serve important mechanical and evolutionary roles (3, 4). The choice of the sister or non-sister chromatid template for repair is thus a key determinant for meiotic recombination outcome.

DNA strand exchange recombinases are central in regulating the choice of DNA template for DSB repair (5, 6). RAD51 and DMC1 recombinases are two eukaryotic RecA homologs and can assemble into nucleofilaments on single-stranded DNA (ssDNA) generated from the processing of DSBs (5, 7). Both recombinases can perform homology searches of the genome and strand invasion on the donor template during meiosis. Cytologically, RAD51 and DMC1 form nuclear foci on the meiotic chromosomes (8–11). Studies in many species argue that meiotic break repair occurs in two temporally distinct phases: A DMC1-permissive phase 1, followed by RAD51-permissive phase 2 (12–17). In the DMC1-permissive phase 1, DMC1 predominantly repairs DSBs and catalyzes interhomolog recombination, whereas RAD51 is kept catalytically inactive (18–22). In the RAD51-permissive phase 2, the RAD51-mediated pathway becomes active to repair remaining DSBs, mainly using sister chromatids (14–17). RAD51-dependent pathway also operates to repair DSB on sisters before the meiotic entry (23).

During phase 1, different RAD51-inhibiting strategies appear to have evolved in eukaryotic species. The Mek1-mediated pathway downregulates Rad51-dependent repair in budding yeast (20, 22). This regulation is, however, not conserved in plants as RAD51 is proficient in repairing breaks in the absence of DMC1, albeit using sister chromatids inferred from a lack of interhomolog COs (24, 25). The mere presence of DMC1 also attenuates the RAD51 repair in both yeast and Arabidopsis (21, 26). Constitutive activation of Rad51, in addition to active Dmc1, elicits a longer repair time in yeast (17). The wild-type level of DMC1-mediated interhomolog recombination nonetheless requires the presence of RAD51 but not its catalytic activity (27, 28). In Arabidopsis, RAD51 fused to GFP at c-terminal (RAD51-GFP) is catalytically inactive to repair breaks in mitotic and meiotic cells but supports DMC1-mediated repair during meiosis (28). This suggested that the catalytic activity of RAD51 is dispensable for DMC1-mediated repair in plants.

*In vivo* functions of DMC1 and RAD51 require many accessory proteins in eukaryotes. In plants, BRCA2 mediates RAD51 and DMC1 foci formation, whereas SDS is specifically required for DMC1 focus formation (29–31). These mediators appear to act *in vivo* at the step of nucleofilament formation. Further, MND1 and HOP2 are evolutionarily conserved proteins required for the DNA exchange activity of DMC1 in plants (32–34). In *mnd1* and *hop2*, DMC1 hyperaccumulation inhibits meiotic DSB repair in Arabidopsis (35–39). However, a weak activity of DMC1 in the *Arabidopsis hop2-2* hypomorphic mutant greatly compromises interhomolog repair and allows the RAD51-dependent DSB repair on sisters (34). RAD54 is also required for RAD51 functions and is dispensable for meiotic DSB repair in the presence of DMC1 in Arabidopsis (40). Further, Arabidopsis has five structurally related RAD51 paralogs: RAD51C, XRCC3, RAD51D, RAD51B, and XRCC2 (41). These paralogs can form a tetrameric complex called BCDX2 complex (42). While Arabidopsis RAD51C and XRCC3 are essential for RAD51 focus formation and meiotic repair (37, 43, 44), RAD51B, RAD51D, and XRCC2 play no critical roles in the RAD51-dependent meiotic DSB repair, irrespective of the presence or absence of DMC1 (40, 41). However, the loss of Arabidopsis RAD51B and XRCC2 slightly increases the meiotic recombination rate, implying their unclear roles in meiotic repair (45).

Meiotic chromosome axis proteins ensure DSB repair and CO formation between homologs in a process called interhomolog bias (46). Arabidopsis ASY1, ASY3, and ASY4 are three-meiotic axis-associated proteins that promote synapsis, a process allowing tethering between homologs through polymerization of synaptonemal complex (SC) proteins such as ZYP1 (47–51). ASY1 localizes on the meiotic axis in an ASY3-dependent manner and is depleted from synapsed regions, following the synaptonemal complex assembly between homologs (50). Loss of ASY1, ASY3, and ASY4 results in a substantial reduction of interhomolog COs, albeit in different magnitudes, with meiotic DSBs predominantly repaired on sisters. ASY1 is required for DMC1 stabilization, suggesting a functional relationship between the meiotic axis and repair machinery (49). How meiotic chromosome axis proteins promote DSB repair between homologs is currently unclear.

Most eukaryotes have two classes of COs formed between homologs. In Arabidopsis, class I constitute a significant proportion (85-90%) of COs, mediated by the ZMM group of proteins (SHOC1, PTD, HEI10, ZIP4, MSH4/5, and MER3) and MLH1/3 (52). Class I CO pathway ensures the obligate CO between homologs but is sensitive to CO inference that avoids class I CO formation near each other (53, 54). Class II COs are derived from the structure-specific endonuclease-dependent pathway, including MUS81 (2). Three non-redundant anti-Class II CO pathways implicating FANCM, RECQ4A & RECQ4B, and FIDGETIN-LIKE-1 (FIGL1) limit meiotic COs by distinct mechanisms (55–60). Although these three pathways regulate class II CO, FIGL1 can also control the distribution of class I COs among chromosomes (60, 61). Loss of Arabidopsis FIGL1 displays a moderate increase in COs with occasional achiasmatic chromosomes (57, 60).

FIGL1 has enigmatic roles in positively and negatively regulating meiotic CO formation. Arabidopsis FIGL1 limits MUS81-dependent COs potentially arising from the resolution of aberrant recombination intermediates, owing to the deregulation of RAD51- and DMC1-mediated strand invasions (57, 60, 62). FIGL1 and its mammalian ortholog FIGNL1physically interact with both recombinases and can antagonize positive mediators of RAD51/DMC1, such as BRCA2 and SDS in Arabidopsis or SWSAP1 in humans (57, 60, 62, 63). *Arabidopsis figl1*, rice *figl1,* and mice *fignl1^cko^*mutants show a change in dynamics of RAD51 and DMC1 foci, arguing for the conserved role of FIGL1/FIGNL1 in meiotic DSB repair (60, 64–67). FIGNL1 is also involved in DSB repair via homologous recombination in somatic cells (63, 68). FIGL1/FIGNL1 is thus a conserved regulator of RAD51- and DMC1-mediated strand invasion and could function in the fine-regulation of strand invasion step to promote accurate meiotic DSB repair. How FIGL1 regulates meiotic break repair outcome when RAD51 and/or DMC1-dependent pathways are fully or partially impaired remains unknown. In this study, we investigated the impact of the functional relationship of *FIGL1* with components of HR repair machinery and chromosome axis genes on the outcome of meiotic DSB repair. We demonstrate that these genetic interactions are an essential determinant of meiotic break repair outcome.

## Results

### 1) Fertility analysis of wildtype and *figl1* plants

We generated ten genotype combinations with *A. thaliana figl1* mutation to study genetic interactions of *FIGL1* with components of meiotic repair machinery and chromosome axis. In all combinations, wild type, single or multiple mutant sister plants grown together were analyzed. When compared to its wild type controls, *figl1* mutants showed slight reduction in fertility (seed set per fruit per plant) that is consistent to previous report (69) but was not always statistically significant (Fig.1, Supp_Data1_SeedCount). The fertility reduction in *figl1* is correlated with a mild defect in meiotic chromosome segregation due to shortage of bivalents in 7% of metaphase I cells as already described in (60). The remaining genotypes are presented in their respective sections.

**Figure 1.**
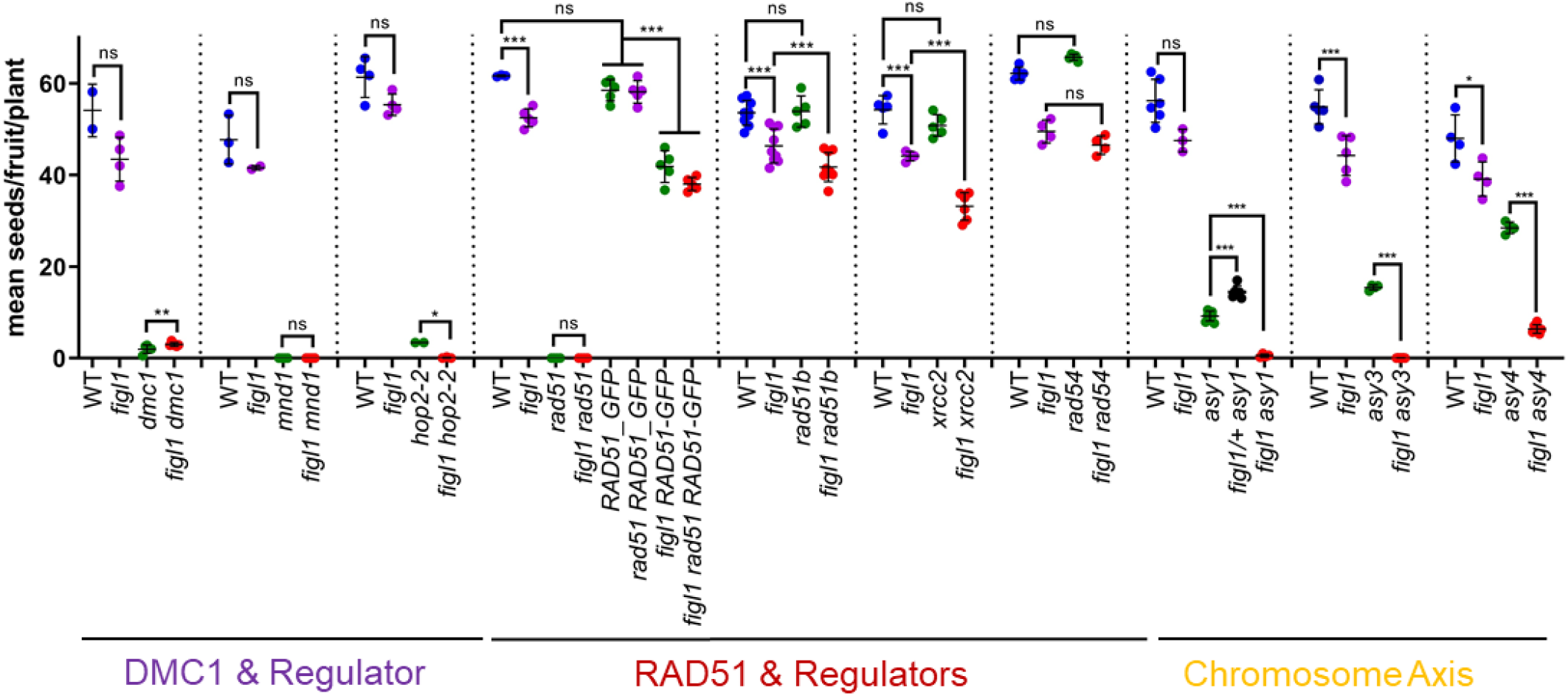
Fertility comparison of *figl1* mutant combined with mutations in genes regulating DMC1/RAD51 or the meiotic chromosome axis in *A. thaliana*. Each colored dot represents one plant with the average number of seeds per fruit counted from at least 10 fruits. The mean and the standard deviation are represented by the black bars for each genotype. Each *figl1* double mutant combination is compared with wild-type (WT) sister plants and respective single mutants that were cultivated together in a segregating population. The P values shown were computed using an unpaired Kruskal-Wallis test corrected with Dunn’s test for multiple comparisons. Ns; non-significant. * indicate a P-value <0.01, ** indicate a P-value <0.001, *** indicate a P-value <0.0001 and ns indicates P > 0.01.

### 2) FIGL1 is dispensable for the RAD51-dependent repair on sisters in *dmc1* and haploids

RAD51 mediates the meiotic DSB repair on sister chromatids in Arabidopsis *dmc1* (24). We explored whether FIGL1 modulates RAD51-mediated meiotic repair without DMC1 by analyzing fertility and male meiosis progression in *figl1 dmc1* double mutant. The *figl1 dmc1* double mutant was sterile with a slightly higher seed set (3 seeds per fruit, plant = 6, fruit=178, corrected Dunn’s test *P* = 0.001) compared to *dmc1* (2 seeds per fruit, plant= 5, fruit=160) (Fig. 1), showing that *figl1* mutation does not restore fertility but can produce bit more seeds in the *dmc1* background. Chromosome spreads of male meiocytes revealed that meiotic progression in *figl1 dmc1* was comparable to *dmc1.* We detected no pachytene stage with fully coaligned homologous chromosomes in *figl1 dmc1*, similar to *dmc1* (Fig. S1). *dmc1* displayed 10 univalents (homologs without CO) at metaphase I cells (N=71) (Fig. 2A), while *figl1 dmc1* exhibited 10 univalents only in 96% of metaphases (N=70). A lack of chromosome fragmentation at meiosis II indicated DSB repair completion in *figl1 dmc1* (Fig. S1). The remaining 4% metaphases (N=3) in *figl1 dmc1* indicated the presence of at least one bivalent (Fig. 2A&2C), that could arise from non-homologous association between homologs or from CO formation between homologous chromosomes. However, we disfavor the former possibility as such non-specific associations would not enhance the fertility in *figl1 dmc1* and are absent in *dmc1*. Thus, loss of FIGL1 neither impairs repair on sisters nor restores CO between homologs in *dmc1*, but FIGL1 appears to counteract rare RAD51-mediated interhomolog repair when lacking DMC1.

**Figure 2.**
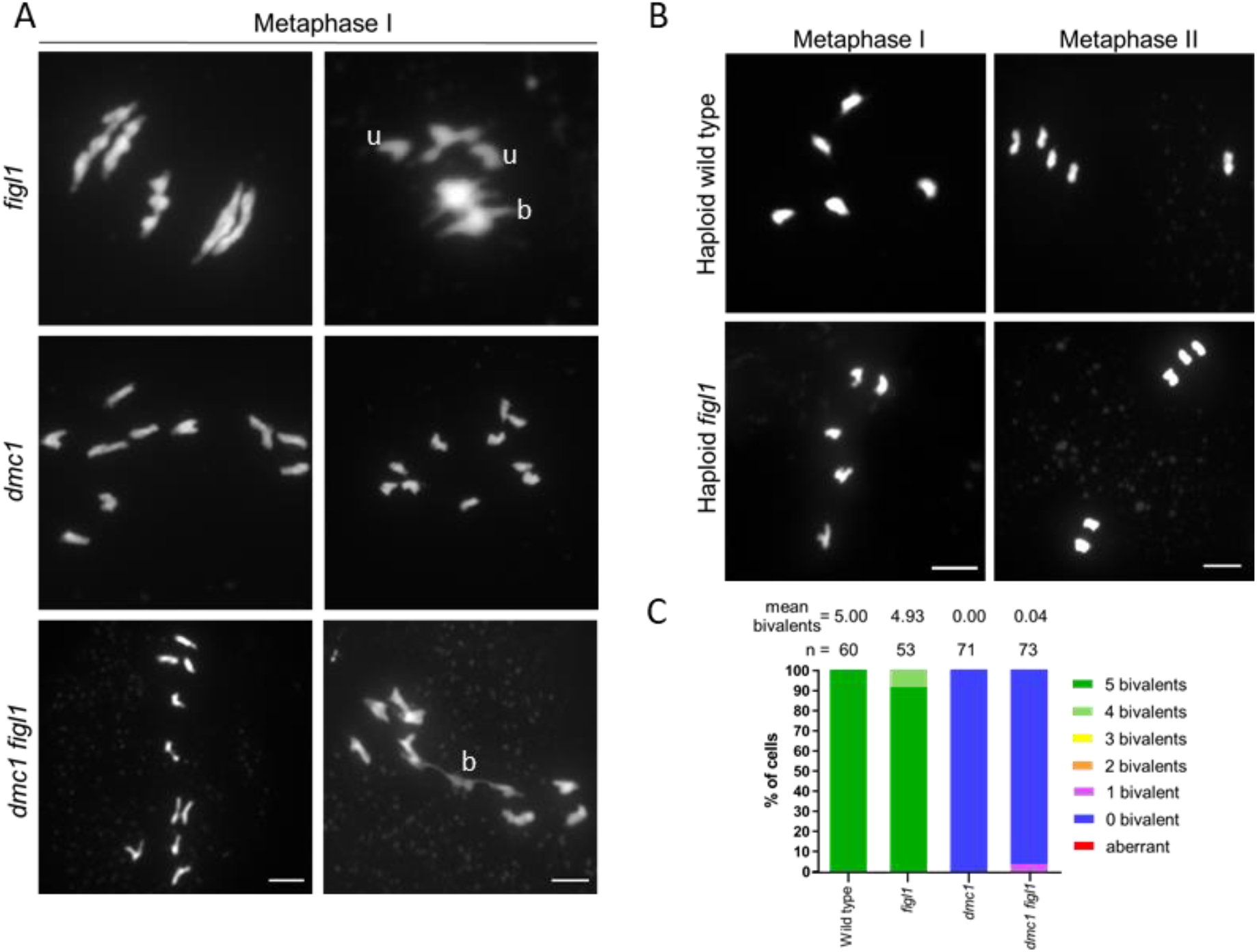
Chromosome spreads display the completion of meiotic break repair in *figl1, figl1 dmc1*, and haploid *figl1* mutant plants. (A) Two representative metaphase I images of DAPI-stained chromosome spread of male meiocytes are shown in *figl1*, *dmc1,* and *figl1 dmc1*; u denotes univalent, while b indicates bivalent or bivalent-like; scale bars: 5µm. B) Metaphase I and II images of DAPI stained chromosome spread from haploid wild-type and *figl1* plants; scale bars: 5µm. C) Quantification of bivalents at metaphase I. The histogram shows the proportion of cells categorized based on the presence of bivalents. The average number of bivalents per cell and the number of analyzed cells are indicated above each bar.

Repairing meiotic breaks relies on RAD51 but not on DMC1 in Arabidopsis haploid meiocytes (70), which lack homologs, and the only available repair template is sister chromatid. We examined if FIGL1 could regulate RAD51-dependent repair on sisters in haploid *figl1*. We analyzed male meiotic progression in Arabidopsis haploid wild-type and *figl1* plants by chromosome spreads. In wild-type haploid meiosis, five univalent chromosomes were intact at metaphase I, but segregated unequally at anaphase I and in variable partitioning later in meiosis II (Fig. 2B). This suggested an efficient repair of meiotic breaks using sister chromatids. No chromosome fragmentation was observed in the *figl1* haploid meiocytes, and meiosis appeared indistinguishable from wild type (Fig. 2B). Altogether, these results suggest that FIGL1 is not required for RAD51-dependent meiotic DSB repair in haploids or *dmc1* but can suppress rare RAD51-mediated interhomolog invasions.

### 3) Downregulation of RAD51-mediated functions in *figl1* alters meiotic DSB repair outcome

We next asked whether RAD51-dependent pathway is dispensable for meiotic repair in the absence of FIGL1. Arabidopsis RAD51-GFP is catalytically inactive and is not required for DMC1-mediated repair (27, 28). We combined *figl1* with *rad51 RAD51-GFP* to evaluate if FIGL1 promotes accurate DMC1-mediated repair in the presence of catalytically inactive RAD51. The fertility of *figl1 rad51 RAD51-GFP* plants (38 seeds per fruit, plants=5, fruit=93, corrected Dunn’s test *P* < 0.0001) was significantly reduced compared to *figl1* (52 seeds per fruit, plant=5, fruit=94) and *rad51 RAD51-GFP* (58 seeds per fruit, plant=5, fruit=90), (Fig. 1, Supp_Data1_SeedCount) but was similar to *figl1 RAD51-GFP* plants (41 seeds per fruit, plant=5, fruit=93, corrected Dunn’s test *P* > 0.99) (Fig. 1). This is consistent with the previous observations of *RAD51-GFP* being dominant-negative (28). These data demonstrate that *figl1* significantly reduces fertility in the *RAD51-GFP* background.

Cytological analysis of meiotic spreads from male meiocytes revealed an elevated frequency of metaphase I cells (37%) with bivalent shortage in *figl1 rad51 RAD51-GFP* (mean bivalents = 4.46, N= 58, *P* =0.0001) compared to 7% cells in *figl1* (mean bivalents = 4.9, N=53) (Fig. 3, Supp_Data2_BivalentCount). The presence of less bivalent in *figl1 rad51 RAD51-GFP* could result from a partial defect in obligate crossover implementation when both FIGL1 and the catalytically active RAD51 are lacking. These defects prompted us to examine class I CO distribution and frequency by immunolocalizing HEI10 and MLH1 at late prophase I in male meiocytes as previously described (71). A similar number of HEI10-MLH1 cofoci was observed in *figl1 rad51 RAD51-GFP* (mean=10.3, n=189, corrected Dunn’s test *P* > 0.99) compared to sister wild-type plants (mean=10, N=75), *figl1* (mean=9.9, N=37) and *rad51 RAD51-GFP* (mean=10.4, N=75) (Fig. 3). The bivalent shortage with wild-type level of MLH1 foci could imply either more intersister repair or defects in CO distribution in *figl1 rad51 RAD51-GFP* compared with *figl1*. In summary, *figl1* does not modulate the number of HEI10-MLH1 foci but significantly reduces bivalent formation when RAD51 is catalytically inactive.

**Figure 3.**
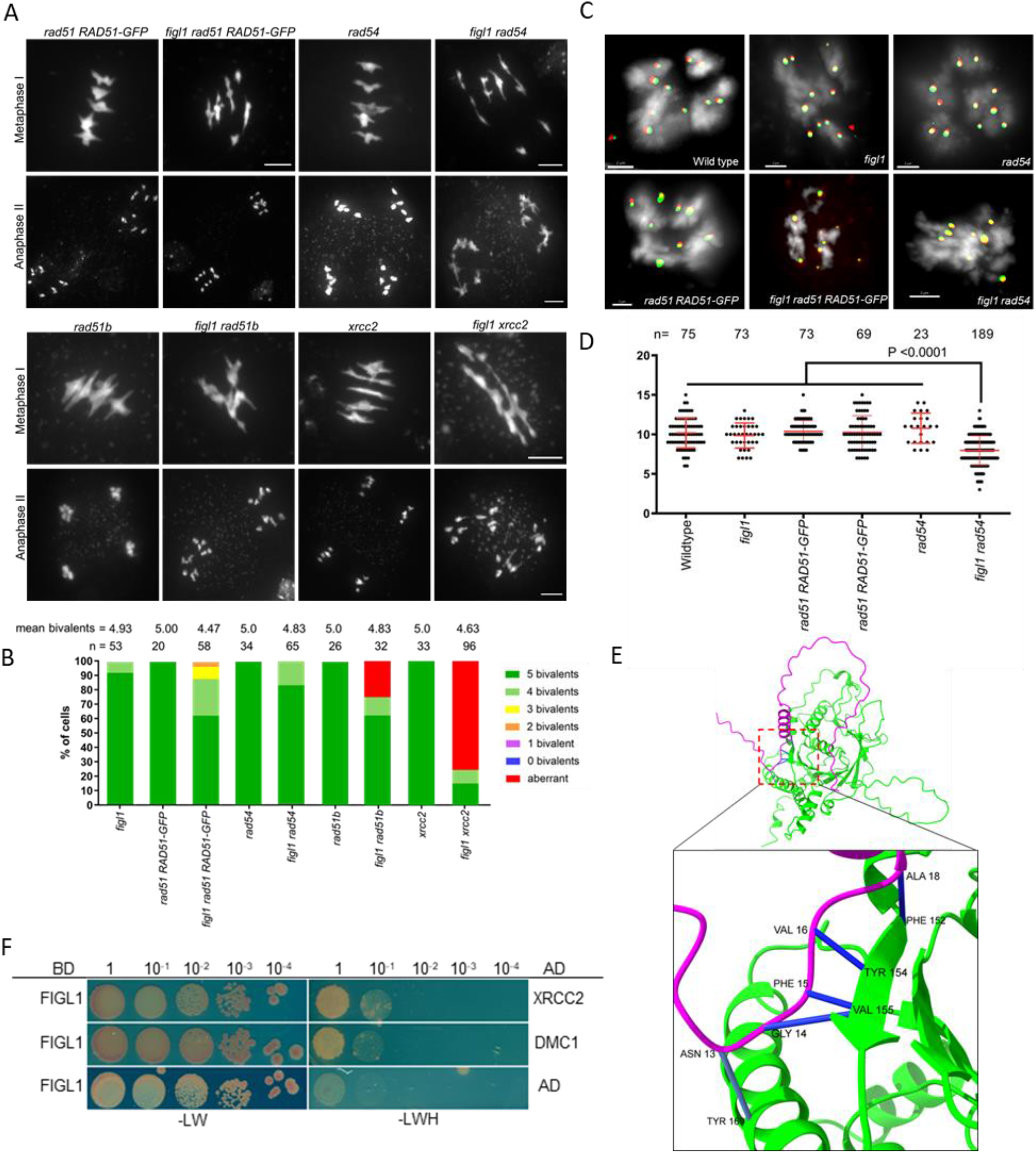
Downregulation of the RAD51-dependent pathway without FIGL1 alters meiotic break repair outcomes. A) Representative metaphase I and anaphase II images of DAPI-stained chromosome spread of male meiocytes are shown in *rad51b*, *figl1 rad51b, xrcc2*, *figl1 xrcc2*, *rad51 RAD51-GFP*, *figl1 rad51 RAD51-GFP*, *rad54*, and *figl1 rad54*; scale bars: 5µm. B) Quantification of bivalents and aberrant repair at metaphase I. The histograms show the percentage of metaphase cells exhibiting bivalents and chromosome fragmentation. The number of analyzed cells (n) and the mean bivalents per cell are indicated above each bar. C) The number of HEI10-MLH1 foci is reduced in *figl1 rad54* but not in *figl1 rad51 RAD51-GFP.* Representative images of HEI10 and MLH1 colocalization in wild type, *figl1, rad54*, *rad51 RAD51-GFP*, *figl1 rad51 RAD51-GFP,* and *figl1 rad54*. scale bars: 2µm. D) Quantification of HEI10-MLH1 cofoci. Each dot represents HEI10-MLH1 cofocus in an individual cell, and the red bars represent the mean per genotype. n = number of cells analyzed. The P values shown were calculated with unpaired Kruskal-Wallis test corrected with Dunn’s test for multiple comparisons. E) 3D structure model of interaction between Arabidopsis XRCC2 (in green) and FIGL-FRBD domain (in magenta) by Alphafold2 structural analysis. An enlarged 3D view of the binding interface between XRCC2 and FIGL1-FRBD domain shows implicated residues. The dark and light Blue lines indicate the high confidence predicted aligned error (PAE) scores for residues with <8Å distances by Alphafold2. F) Yeast-two-hybrid assays showing the interaction of Arabidopsis FIGL1 with XRCC2 and DMC1 (used as positive control). Proteins of interest were fused with the Gal4 DNA binding domain (BD, left) and with the Gal4 activation domain (AD, right), respectively, and co-expressed in yeast cells with selection on SD/-LW (- Leu -Trp) for diploid strains and on SD/-LWH (- Leu -Trp -His) for protein interaction.

Arabidopsis *rad54* mutants show no meiotic defects but display massive chromosome fragmentation in the *rad54 dmc1* (40), indicating an essential requirement of RAD54 for RAD51-mediated repair but not for DMC1-dependent repair. We next tested the epistatic relationship between *rad54* and *figl1.* Fertility of *rad54* (65 seeds per fruit, plant=5, fruit=80) compared to wild type (62 seeds per fruit, plant=5, fruit=82) was not affected as described in (40). The fertility analysis of *figl1 rad54* (46 seeds per fruit, plant=4, fruit=70) showed a slightly lower seed set, which is not significantly different from *figl1* (49 seeds per fruit, plant=4, fruit=67, corrected Dunn’s test *P* =0.57) (Fig. 1). Meiotic chromosome spreads analysis revealed that *rad54* and the wild type displayed 100% metaphase I cells with five bivalents but *figl1 rad54* showed 16.9% of metaphases with bivalent shortage compared to 7% in *figl1.* The mean bivalent per cell in *figl1 rad54* (4.8, N=65) was not significantly different from *figl1* (4.9, N=53).

However, the mean HEI10-MLH1 cofocus formation at late prophase I was ∼20% reduced in *figl1 rad54* (8 foci, N=189, corrected Dunn’s test *P <0.0001*) compared to *figl1* (9.9 foci, N=37), *rad54* (11 foci, N=23), and wild type (10 foci, N=75) (Fig. 3). These data suggested that the loss of RAD54 activity aggravates phenotypes in *figl1* with reduced number of HEI10-MLH1 foci and increased number of cells with less bivalents. Taken together, our data indicate that lack of FIGL1 elicits a requirement of the RAD51-dependent pathway to ensure the wild-type level of meiotic COs or chiasmata. This could also imply that FIGL1 likely suppresses RAD51-dependent repair pathway during the wild-type meiotic repair.

### 4) XRCC2 and RAD51B are required for the repair of the meiotic breaks in *figl1*

Lack of RAD51 strongly perturbs DMC1-dependent repair (72). We next examined if the deficiency of both FIGL1 and RAD51 could facilitate DMC1-mediated repair by generating the *figl1 rad51* double mutant. Both *rad51* and *figl1 rad51* showed complete sterility with the mean seed set per fruit of zero (*rad51:* 0 seeds per fruit, plant= 2, fruit=56; *figl1rad51:* 0 seeds per fruit, plant=3, fruit=70) (Fig. 1, Supp_Data1_SeedCount). Thus, *rad51* sterility is unchanged in the *figl1* background. Male meiotic progression was also indistinguishable in *rad51* and *figl1 rad51,* with strong chromosome fragmentation indicating unrepaired meiotic breaks (Fig. S2). These observations suggest that *rad51* is epistatic to *figl1* and that loss of FIGL1 and RAD51 together did not facilitate meiotic repair.

The cellular activity of RAD51 is regulated by the RAD51 paralogs, including RAD51B and XRCC2 (41). We examined whether *RAD51B* and *XRCC2* genetically interacted with *FIGL1* by analyzing *figl1 rad51b* and *figl1 xrcc2* double mutants for fertility and meiotic defects. Similar to previous observations (40, 45, 73), the fertility of *rad51b* and *xrcc2* did not differ from their sister wild-type plants (Fig. 1, Supp_Data1_SeedCount). However, both *figl1 rad51b* (42 seeds per fruit, plant=9, fruits=56, corrected Dunn’s test *P <*0.0001) and *figl1 xrcc2* (33 seeds per fruit, plant =6, fruit=44, corrected Dunn’s test *P <*0.0001) mutants showed reduced fertility compared to *figl1*, and respective *rad51b* and *xrcc2* (Fig. 1). Thus, loss of FIGL1 affects fertility differentially in *rad51b* and *xrcc2* backgrounds. Our cytological analysis of male meiosis supported a different severity level for meiotic defects. As previously reported, no visible meiotic defects were detected in *rad51b* and *xrcc2* mutants (40, 45). The *figl1 rad51b* displayed 25% metaphases I with aberrant chromosome structures and 10 % metaphases with a mixture of univalent and bivalents, suggesting a defect in meiotic repair (Fig. 3). The *figl1 xrcc2* displayed more severe defects as 75% of metaphases I had chromosome fragmentation, univalents, multivalents, or aberrant structures (Fig. 3). These meiotic catastrophes in both *figl1 rad51b* and *figl1 xrcc2* were also confirmed in subsequent meiotic stages leading to unbalanced chromosome segregations (Fig. 3A). Altogether, *rad51b* and *xrcc2* do not exhibit any meiotic defects but significantly impair DSB repair in the *figl1* context, revealing that their functional interaction with *FIGL1* promotes meiotic DSB repair.

### 5) FIGL1 suppresses partial defect of RAD51 focus formation in *xrcc2* meiocytes

Our data suggested that Arabidopsis XRCC2 and FIGL1 could act as positive and negative regulators of RAD51 assembly, respectively. We next explored if the functional relationship between *XRCC2* and *FIGL1* could regulate RAD51 assembly in male meiocytes by immunolocalizing RAD51 and axial element (REC8). RAD51 focus formation was observed in wild-type, *xrcc2*, *figl1*, and *figl1 xrcc2* meiocytes during prophase I stages, identified by REC8 signal along the chromosome axis (Fig. 4A). The mean RAD51 focus number was 27% reduced in *xrcc2* (101 foci, N=75, *P* <0.0001) compared with wild-type meiocytes (138 foci, N=103) (Fig.4B). Hence, XRCC2 is required for the formation and/or stabilization of a subset of RAD51 foci during prophase I. *figl1* mutation suppressed this partial defect in RAD51 focus formation in *xrcc2*. The *figl1 xrcc2* (254 foci, N=75, *P* <0.0001) meiocytes showed a significantly higher number of RAD51 foci compared to *xrcc2* and wild type (138 foci, N=103) but similar to *figl1* (204 foci, N=89) (Fig. 4B). These data suggest that FIGL1 could disrupt a subset of RAD51 foci in *xrcc2* and that XRCC2 would protect RAD51 filaments by partially antagonizing FIGL1 activity during meiosis.

**Figure 4.**
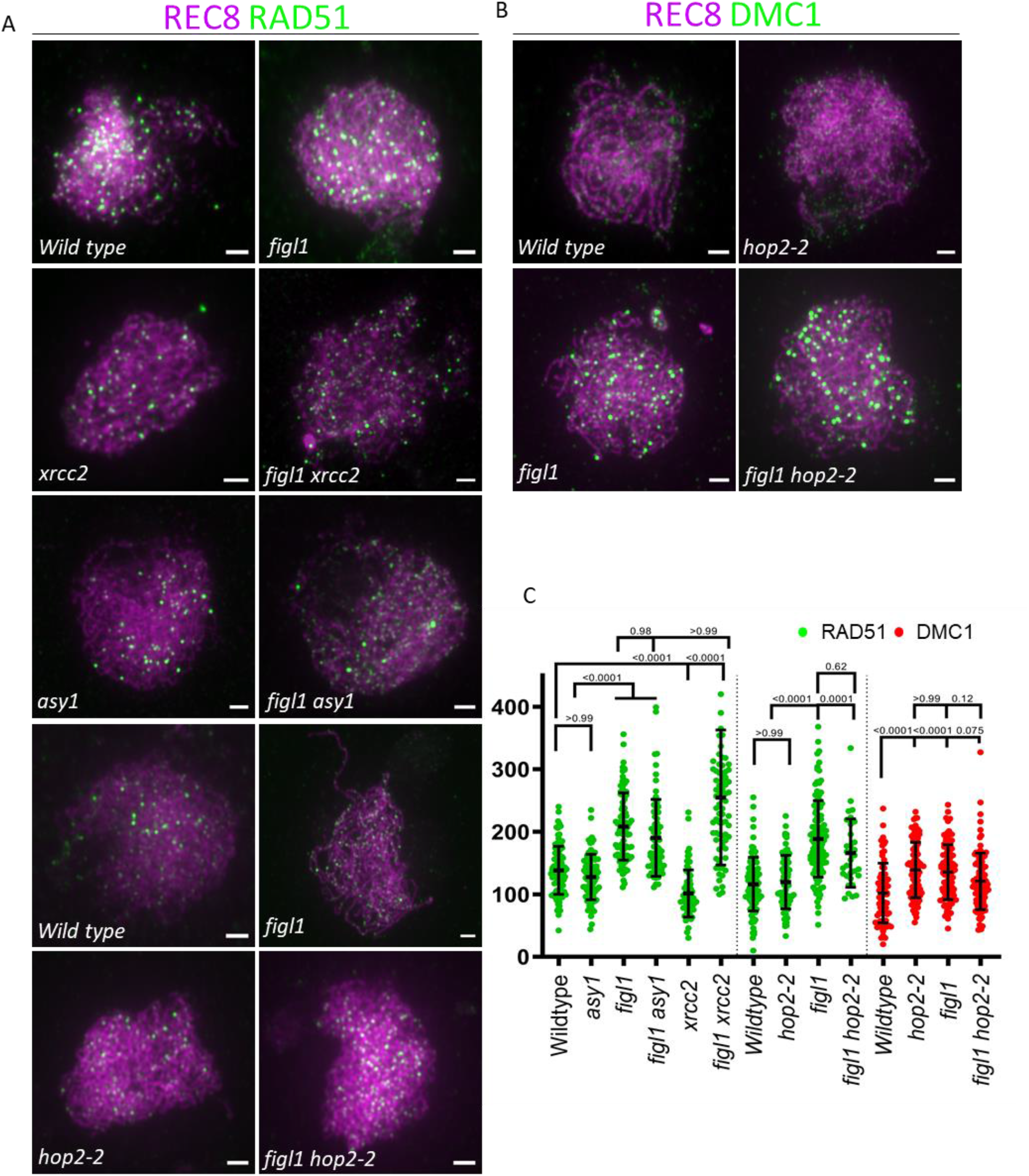
Analysis of distribution of RAD51 or DMC1 in different *figl1* mutants. A) Dual immunolocalization of REC8 (magenta) and RAD51 (green) on male meiocytes from wild type (Col- 0), *figl1*, *xrcc2*, *figl1 xrcc2*, *asy1*, *figl1 asy1*, and wild type, *figl1, hop2-2*, *figl1 hop2-2* in the Col-0/Ws hybrid background. B) Dual immunolocalization of REC8 (magenta) and DMC1 (green) o male meiocytes from wild type, *figl1, hop2-2*, and *figl1 hop2-2* in the Col-0/Ws hybrid background. C) Quantification of RAD51 and DMC1 foci in different wild types and mutants. The P values shown were calculated with unpaired Kruskal-Wallis test corrected with Dunn’s test for multiple comparisons.

In humans, RAD51 paralogs can protect RAD51 filaments by antagonizing the FIGL1 homolog through protein-protein interaction (63). We therefore asked if Arabidopsis XRCC2 and FIGL1 could also interact. Since XRCC2 has structural similarities with RAD51 that interacts with FIGL1 through FIDGETIN-LIKE-1’s RAD51 Binding Domain (FRBD) (60, 68), we generated an interaction model of XRCC2 and FRBD from FIGL1 by Alphafold2. All five Alphafold2 models predicted a strong interaction with a high interface-predicted Template Modeling score (ipTM) (>0.60) and a low predicted aligned error (PAE) value for the FxxA motif in FRBD and three residues of XRCC2 (F152, W154 & V155) (Fig. 3E and S3). We also tested interaction of RAD51B and FRBD by Alphafold2, which did not predict a high ipTM score (<0.3) compared to XRCC2 and FRBD interaction (Fig. S3). These observations suggest that XRCC2 likely interact with FIGL1 through FRBD, more strongly than RAD51B and that this could be reminiscent of FIGL1 and RAD51 interaction. Yeast two-hybrid assay indeed confirmed the interaction between full-length Arabidopsis XRCC2 and FIGL1 (Fig. 3F). These results demonstrate an interaction between XRCC2 and FIGL1 and support the hypothesis that XRCC2 counteracts FIGL1 to protect a subset of RAD51-filaments.

### 6) FIGL1 is indispensable for repair completion in *hop2-2*

DMC1 requires MND1 to facilitate strand invasion on homologs (35, 36). We wondered if FIGL1 modulates meiotic repair defect when DMC1 cannot perform interhomolog invasions in the *mnd1* background by generating the *mnd1 figl1* mutant. Both *mnd1* (0 seeds per fruit, plant=4, fruit=108) and *mnd1 figl1* (0 seeds per fruit, plant = 5, fruit=86) mutants are fully sterile with empty siliques (Fig. 1), suggesting that *figl1* does not change the fertility of *mnd1*. Similar to previous observations (35), analysis of male *mnd1* meiocyte spreads revealed no pachytene cells with synapsed chromosomes and the presence of a strong chromosome fragmentation at meiosis I and II (Fig. 5A). This supports a lack of interhomolog strand invasion and unrepaired breaks in *mnd1*. The double *mnd1 figl1* also showed meiotic progression with chromosome fragmentation, which was indistinguishable from *mnd1* (Fig. 5A), showing that *mnd1* is epistatic to *figl1*.

**Figure 5.**
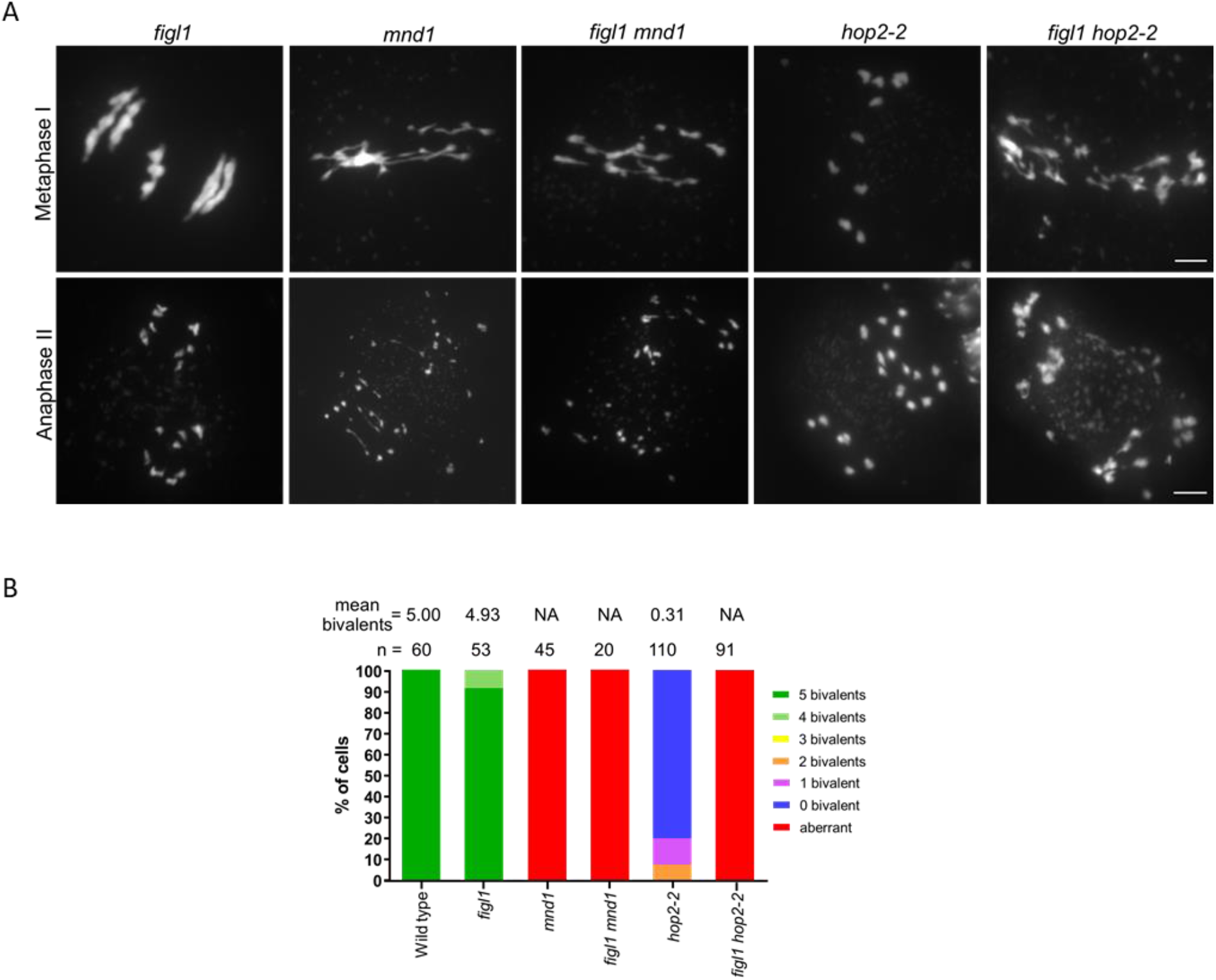
Epistatic analysis of *mnd1*, *hop2-2*, and *figl1*. (A) Representative metaphase I and anaphase II images of DAPI-stained chromosome spread of male meiocytes are shown in *figl1*, *mnd1, hop2-2*, *figl1 mnd1*, and *figl1 hop2-2*; scale bars: 5µm. B) Quantification of bivalents and aberrant repair at metaphase I. The histogram shows the proportion of metaphase cells categorized based on the presence of bivalents and chromosome fragmentation. The mean bivalents per cell and the number of analyzed cells (N) are indicated above each bar. NA; not applied

A *hop2-2* hypomorphic mutation weakly supports DMC1 functions that lead to the RAD51-dependent DSB repair on sisters and severely perturb interhomolog repair (34). We investigated if FIGL1 is implicated in meiotic repair when DMC1 is not fully active by analyzing *figl1 hop2-2* double mutant. The fertility of *figl1 hop2-2* plants (0.1 seeds per fruit, plants = 3, fruit=64) was reduced compared to *hop2-2* (3 seeds per fruit, plant = 2, fruit=40), wild type (62 seeds per fruit, plant= 4, fruit=76), and *figl1* (54 seeds per fruit, plant= 3, fruit=75) (Fig. 1, Supp_Data1_SeedCount). Cytological analysis of metaphase I from male *figl1 hop2-2* meiocytes further corroborated the reduction of fertility with defects in meiotic progression. As reported previously (34), we detected only 20% of metaphase I with 1 to 3 bivalents and the rest with only univalent chromosomes in the male *hop2-2* meiocytes (Fig. 5b). In the *figl1 hop2-2*, all metaphase I (N=91) and subsequent meiotic stages exhibited strong chromosome fragmentation compared to the *hop2-2* meiocytes (Fig. 5). These results indicate that *figl1* strongly perturbs meiotic repair in *hop2-2* and implicate FIGL1 in meiotic repair switching from interhomolog to mostly intersister when DMC1 is partially active.

We next examined if meiotic repair defects in *figl1 hop2-2* could be explained by hyperaccumulation of both recombinases during prophase I by immunolocalizing RAD51 or DMC1 with the axial element (REC8) and the transverse element of the SC (ZYP1) in male meiocytes. Average RAD51 foci per cell during prophase I did not differ between wild-type (foci= 115, N= 104) and *hop2-2* (foci= 114, N= 76) meiocytes but were significantly higher in *figl1 hop2-2* (foci= 155, N= 31) and *figl1* (foci= 186, N= 116). A marked increase in mean DMC1 foci per cell was observed in *hop2-2* (foci= 138, N= 93) meiocytes relative to wild type (foci= 101, N= 74), but it was similar to *figl1* (foci= 135, N= 93) and *figl1 hop2-2* (foci= 120, N= 100) (Fig. 4). These observations suggest that FIGL1 does not change the number of DMC1 foci but appears to suppress RAD51 focus formation in *hop2-2*. We have previously shown that *figl1* mutation restores RAD51 and DMC1 focus formation and suppresses defects in synapsis between homologs in *brca2a brca2b* and *sds* (60, 62). Since no pachytene is observed in *hop2-2* suggesting a defective synapsis, we investigated if synapsis was restored partially or fully in *figl1 hop2-*2 by ZYP1 immunostaining. Only punctuated ZYP1 signal was observed in both *figl1 hop2-2* relative to *hop2-2*, while wild-type and *figl1* meiocytes showed continuous ZYP1 signal along the entire length of chromosomes (Fig. S4). These results imply that over-accumulation of RAD51 and DMC1 foci does not restore interhomolog invasion in *figl1 hop2-2* and that FIGL1 likely promotes meiotic repair in *hop2-2* by suppressing RAD51 foci.

### 7) FIGL1 promotes meiotic repair in *asy1*, asy3, and *asy4* with impaired interhomolog bias

We further explored whether FIGL1 is critical for meiotic repair when both RAD51 and DMC1 are active but interhomolog bias is impaired. Unlike in *hop2-2,* DMC1 is proficient for meiotic repair in *asy1, asy3,* and *asy4* mutants, which display a lack of full synapsis with no pachytene stage and a shortage of bivalent formation (49–51). *asy1* is epistatic over *asy3* and *asy4*, and shows the lowest number of bivalents at metaphase I followed by *asy3* and *asy4* (50, 51). The reduction in interhomolog CO, no fragmentation and the bivalent shortage in *asy* mutants suggest that most meiotic DSBs are efficiently repaired on sisters instead of homologs in the wild type. We generated *figl1 asy1*, *figl1 asy3*, and *figl1 asy4* double mutants and analyzed fertility and meiotic defects. All three, *asy1* (10 seeds per fruit, plant=2, fruit=23), *asy3* (15 seeds per fruit, plant=3, fruit=83), and *asy4* (28 seeds per fruit, plant=4, fruit=80) showed a severe reduction in fertility, albeit to a variable extent, compared to *figl1* (44 seeds per fruit, plants=5, fruit=100) and wild type (54 seeds per fruit, plants=5, fruit=100). Both *figl1 asy1* and *figl1 asy3* mutants were sterile and barely produced any seeds (*figl1 asy1*: 0.6 seeds per fruit, plant=4, fruit=70, corrected Dunn’s test *P <*0.0001; *figl1 asy3*: 0 seeds per fruit, plant=7, fruit=138, corrected Dunn’s test *P <*0.0001) (Fig. 1, Supp_Data1_SeedCount). The *figl1 asy4* plants produced a few seeds, but fertility was >4-fold decreased (6 seeds per fruit, plant=7, fruit=140, corrected Dunn’s test *P <*0.0001), compared to *asy4* and *figl1* (Fig. 1, Supp_Data1_SeedCount). Intriguingly, the fertility of *figl1/+ asy1* (14 seeds per fruit, plant=6, fruit=95, corrected Dunn’s test *P <*0.0001) was significantly increased compared with *asy1* (9 seeds per fruit, plant=8, fruit=106) and no significant difference in fertility was observed in *figl1/+ asy3* (20 seeds per fruit, plant=6, fruit=60, corrected Dunn’s test *P* 0.074) and *asy3* (15 seeds per fruit, plant=9, fruit=133), suggesting that meiotic repair is sensitive to FIGL1 dosage in *asy1*. In summary, while the *figl1* mutation affects marginally fertility in the wild-type context, it strongly reduces fertility in *asy1*, *asy3*, and *asy4* mutants of meiotic axis components.

Further, nuclear spread analysis of male meiocytes in *asy1*, *asy3*, and *asy4* showed mean bivalent numbers of 1.66 (N=50), 3.0 (N=67), and 4.1 (N=38) at metaphase I, respectively (Fig. 6). This confirmed the previous observations of *asy1* being most affected in CO formation (49–51). The *figl1 asy1* mutants exhibited 82% of aberrant metaphases (N=52) with unrepaired meiotic breaks, which was never observed in *figl1* and *asy1* (Fig. 6A&B). We, however, detected a fraction of non-aberrant metaphases (18%) without any fragmentation but showing a mixture of univalent and bivalent chromosomes with a higher mean bivalent per cell (3.22, N=10) compared with *asy1*. These results indicate that *figl1* strongly impairs meiotic repair in most *asy1* meiocytes but also partially restores bivalent formation between homologs in a fraction of meiocytes and support that FIGL1 suppresses interhomolog repair in *asy1*. We, therefore, analyzed if the reduction of FIGL1 dosages could increase bivalent formation in *figl1/+ asy1*. A significant increase in mean bivalent per cell (2.5, N=52) at metaphase I was observed in *figl1/+ asy1* compared to *asy1*. These observations further supported that FIGL1 attenuates interhomolog repair when lacking *asy1* and implied that ASY1 counteracts FIGL1 to promote interhomolog repair.

**Figure 6.**
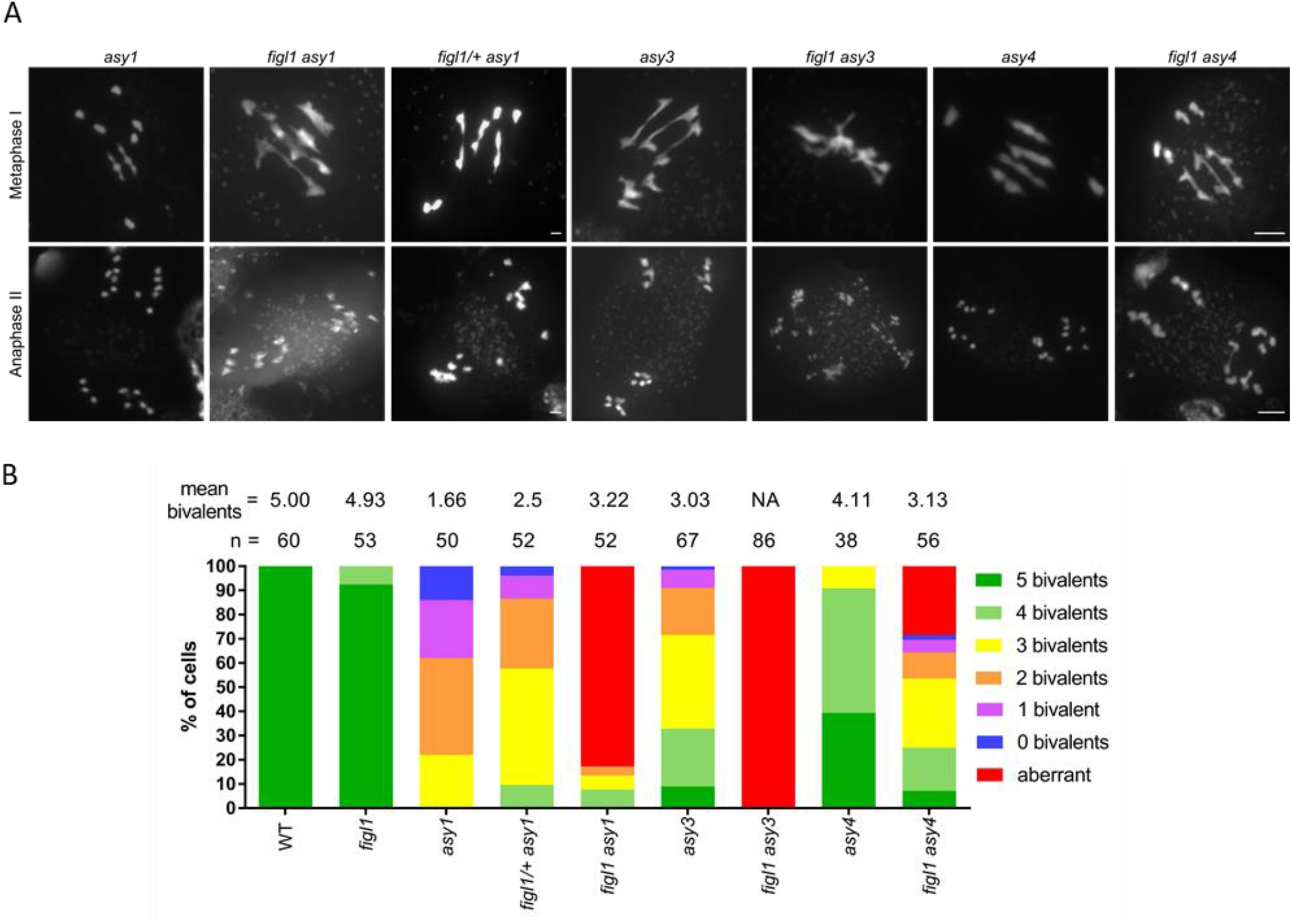
Functional interaction of *figl1* with *asy1*, *asy3*, and *asy4*. (A) Representative metaphase I and anaphase II images of DAPI-stained chromosome spread of male meiocytes are shown in *asy1*, *asy3, asy4*, *figl1 asy1*, *figl1/+ asy1*, *figl1 asy3*, and *figl1 asy4*; scale bars: 5µm. B) Quantification of bivalents and aberrant repair at metaphase I. The proportion of metaphase cells categorized based on the presence of bivalents and chromosome fragmentation is presented. The mean bivalents per cell and the number of analyzed cells (N) are indicated above each bar. NA; not applied

Although *asy3* is less affected than *asy1*, the *figl1 asy3* exhibited the most severe meiotic defect of 100% aberrant metaphases (N=86) with chromosome fragmentation. This indicates that FIGL1 is indispensable for meiotic repair in *asy3* (Fig. 6). The *figl1 asy4* mutants, however, display the least severe meiotic defects among the three *figl1 asy* mutants. 75% of metaphase showed a marked reduction in mean bivalent per cell in *figl1 asy4* (3.13, N=56) compared to *asy4* (4.1, N=38) (Fig. 6). The remaining 25% metaphases in *figl1 asy4* were aberrant with unrepaired breaks (Fig. 6A&B). Meiotic defects in all three *figl1 asy* double mutants were also detected at the post metaphase I stages, showing chromosome fragmentation, bridges, and unbalanced chromosomal segregation (Fig. 6A). In summary, *figl1* severely impairs meiotic repair in the *asy1*, *asy3*, and *asy4* mutant affected in interhomolog bias, and appears to elevate interhomolog repair in *asy1*.

We further assessed whether the repair defects in *figl1 asy1* led to a change in RAD51 focus formation by immunolocalizing RAD51 and REC8 in male meiocytes. Similar to previous observations (49), the number of RAD51 foci was not different in *asy1* (foci=127, N=94) and wild-type meiocytes (foci=138, N=103) (Fig. 4). As expected, both *figl1* (foci=208, N=89) and *figl1 asy1* (foci=190, N=73) showed a significant increase in RAD51 foci compared to *asy1* and wild type, but did not differ from each other. These observations indicate that FIGL1 limits over-accumulation of RAD51 foci in *asy1* to promote meiotic repair. However, the outcomes of RAD51 foci accumulation in *figl1* and *figl1 asy1* differ as repair is fully completed in *figl1* but not in *figl1 asy1*.

## Discussion

### RAD51-dependent repair in *figl1* is critical to ensure correct meiotic repair outcome

Our data indicate a functional association between FIGL1 and RAD51-dependent repair pathway during meiosis. RAD51 pathway is critical to repair DSB before/at meiotic entry and in late prophase I (14, 15, 23), providing an alternative to the DMC1-dependent pathway outside phase 1. However, simultaneous activation of Dmc1 and Rad51 in budding yeast elicits distinct factors and a longer time to repair Rad51-mediated interhomolog intermediates (17). Arabidopsis *figl1* mutants display a change in kinetics with massive accumulation of RAD51 foci on meiotic chromosomes until late pachytene compared with wild-type meiocytes (60). We speculated that RAD51, besides DMC1, is active for interhomolog invasions, leading to aberrant recombination intermediates in *figl1*. These intermediates required MUS81 endonuclease activity that becomes indispensable for repair completion in *figl1 mus81* (57, 62). FIGL1 is, however, essential for meiotic DSB repair in different crop plants, including rice *figl1* mutant showing chromosome fragmentation at metaphase I (64, 66). These unrepaired breaks might result from complex DMC1-and RAD51-mediated interhomolog invasions, which MUS81 is unable to repair, given a more considerable genome complexity and chromosome number in rice (74). Our data provide evidence that the RAD51-dependent pathway promotes accurate meiotic repair outcomes in Arabidopsis *figl1*. First, the inhibition of RAD51’s catalytic activity in *figl1* results in a higher percentage of metaphase cells with univalent, yet with wild-type levels of HEI10-MLH1 foci during prophase I. This suggests either a defect in CO distribution or CO forming between sister chromatids in *figl1 rad51 RAD51-GFP*. Second, downregulation of RAD54 that impairs RAD51-dependent repair is required to maintain the wild-type level of HEI10-MLH1 foci as judged from ∼20% reduction in *figl1 rad54*. Third, the loss of RAD51B or XRCC2 leads to unrepair breaks in *figl1 rad51b* and *figl1 xrcc2*. This indicates that the RAD51-dependent pathway becomes limiting for recombination when lacking FIGL1. Thus, it is tempting to speculate that FIGL1 suppresses RAD51-dependent interhomolog repair to limit class II CO (Fig.7) and regulate class I CO distribution in wild-type meiosis in Arabidopsis.

**Figure 7.**
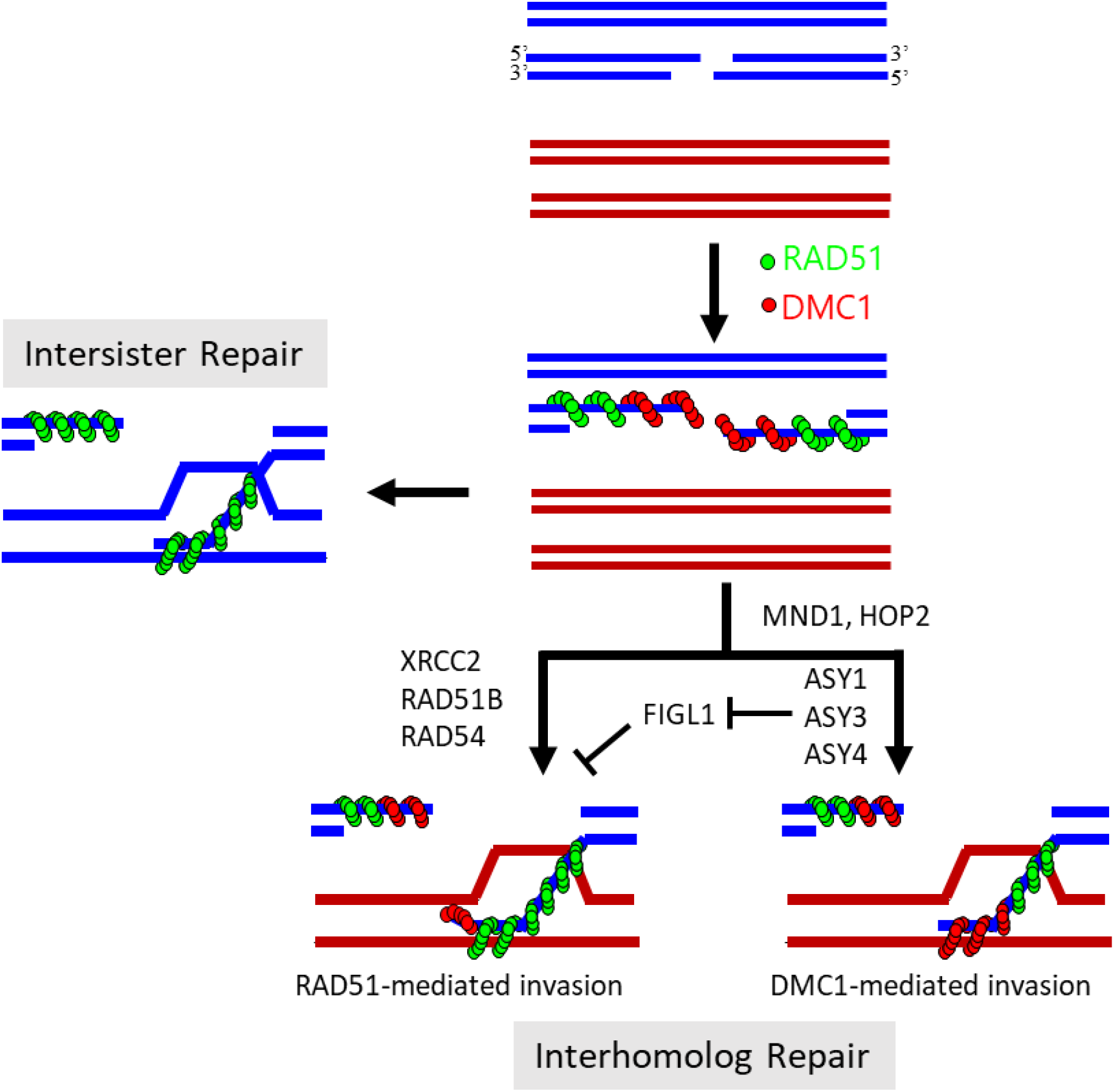
Model for FIGL1 in vivo functions during meiotic break repair. Blue and dark brown colors represent homologous chromosomes, while four lines in the same color denoting two DNA strands of each sister chromatid. After DSB formation and the resection of 5’ ends, RAD51 and DMC1 polymerize at 3’ single-strand DNA (ssDNA) tails to form nucleoprotein homofilaments. DMC1 is proximal to DSB site, while RAD51 is distal to DSB site. Invasion of these homofilaments leads to repair meiotic breaks. FIGL1 can negatively regulate RAD51 and DMC1 activity at pre- and post-invasion steps. FIGL1 is not required for RAD51-mediated intersister repair of meiotic break in absence of DMC1 in plants. However, FIGL1 can antagonize RAD51-mediated interhomolog invasions in presence of DMC1. MND1 and HOP2 likely act upstream of FIGL1. The three meiotic axis proteins (ASY1, ASY3, and ASY4) act as positive modulator of interhomolog recombination by counteracting FIGL1 activity. In absence of FIGL1, RAD51 is likely potent for interhomolog invasions/repair in XRCC2 and RAD51B-dependent manner. The catalytic activity of RAD51 or its stimulation by RAD54 are critical in ensuring the wild-type level or distribution of meiotic CO/chiasmata formation.

### Arabidopsis XRCC2 partially counteracts FIGL1 to promote RAD51-dependent repair

The roles of Arabidopsis RAD51B, RAD51D, and XRCC2 during meiosis remained enigmatic despite the proposed early and late roles of these RAD51-mediators in other plant species (75–77). No obvious meiotic defects in single, double, and triple mutants of Arabidopsis *RAD51B*, *RAD51D,* and *XRCC2* indicate their non-essentiality in DMC1-mediated repair during meiosis (40, 41, 73). Mutants of *rad51b*, *rad51d*, and *xrcc2*, when combined with *dmc1*, have no impact on meiosis, suggesting their dispensability for RAD51-mediated repair on sisters (40). However, an increase in meiotic COs between homologs in Arabidopsis *xrcc2* and *rad51b* remains unexplained (45). Our findings demonstrate a genetic interaction between *xrcc2* and *rad51b* with *figl1*, which plays a critical role in meiotic DSB repair. Firstly, both *figl1 rad51b* and *figl1 xrcc2* mutants have severe meiotic defects, although in varying degrees. This means that both paralogs are necessary for repairing meiotic breaks when FIGL1 is absent, with loss of XRCC2 producing more severe meiotic defects. Recent structural studies of the human BCDX2 complex suggested that RAD51 ensemble interacts with the BCDX2 complex by engaging with RAD51B but not with XRCC2 (78, 79). We speculate that the BCD complex in can still engage with RAD51-filaments through RAD51B, leading to more toxic recombination intermediates and severe meiotic defects in *figl1 xrcc2*, compared to no interaction of CDX2 complex in *rad51b* with RAD51 ensemble. Secondly, we observed a decrease in RAD51 focus number in the *xrcc2* meiocytes compared to the wild type and an increase in RAD51 focus number in *figl1 xrcc2* and *figl1*. This suggests that XRCC2 stabilizes or protects a subset of RAD51 filaments from the destabilizing activity of FIGL1, arguing for a partial antagonism between XRCC2 and FIGL1. We also detected a protein-protein interaction between XRCC2 and FIGL1, which is consistent with the model wherein RAD51 paralogs interaction counteracts destabilizing functions of FIGL1 or other negative regulators (63, 80). We speculate that XRCC2 could counteract FIGL1 activity from the non-engaging end of BCDX2 complex and acts, in addition to BRCA2, at the post nucleofilament assembly step during meiosis.

Furthermore, our data also imply that when both RAD51 and DMC1 are active for homolog invasions (Fig.7), the RAD51-dependent repair pathway has additional requirements compared to those when DMC1 is absent or not functioning. One possibility is that XRCC2 and RAD51B are needed to process a subset of RAD51-mediated invasions, especially in *figl1*, when RAD51 is probably blocked from polymerizing to 3’ ends (Fig.7). XRCC2 can also promote the processing of the recombination intermediates through its interaction with RAD51D to form a subcomplex (DX2) (81), which can recruit additional factors (77, 82). Altogether, our data support that Arabidopsis XRCC2 and RAD51B promote RAD51-dependent repair during meiosis.

### ASY1 counteracts FIGL1 activity to promote interhomolog recombination

Arabidopsis ASY1, ASY3, and ASY4 are three meiotic axis proteins whose mechanistic understanding in interhomolog repair remains elusive. ASY1 appears to be a main player as judged from the lowest chiasma level and epistatic analysis among *asy* mutants (51). The lack of ASY1 does not affect total MLH1 focus numbers, but 50% occur between sister chromatids, indicating reduced interhomolog and elevated intersister recombination levels in *asy1* (83). ASY1 stabilizes DMC1 onto DSBs without affecting RAD51 localization (49). The residual interhomolog recombination in *asy1* still depends on DMC1 as the *asy1 dmc1* mutant shows only univalents with DSBs repaired by RAD51 using sisters (49). Thus, the loss of ASY1 activates both RAD51- and DMC1-dependent pathways to repair meiotic breaks. Our findings show that FIGL1 is required to maintain the balance between interhomolog and intersister recombination to complete the repair in *asy1.* The *figl1 asy1* is completely sterile, and 82% of metaphase I cells exhibit an aberrant phenotype with DNA fragmentation. We noticed that 18% metaphase I cells showed no fragmentation but a significantly higher mean of 3.22 bivalents in *figl1 asy1* compared with 1.66 bivalents in *asy1* (Fig. 6). Furthermore, the reduction of FIGL1 dosages in *figl/+ asy1* also led to an increase in mean bivalent number as well as a higher mean seed set per fruit compared to *asy1* (Fig.1&6). This indicates that FIGL1 suppresses interhomolog invasion in the absence of ASY1, and the repair defects in *figl1 asy1* could thus be attributed to an increase in interhomolog invasions. Altogether, ASY1 counteracts FIGL1 activity to promote interhomolog recombination.

Our data also indicate a functional relationship between *asy3, asy4*, and *figl1* for the completion of meiotic break repair. The double mutant *figl1 asy3* exhibited the most severe repair defect phenotype due to the presence of chromosome fragmentation in 100% of metaphase I (Fig. 6). Lack of ASY3 significantly changes localization of ASY1 that forming foci instead of linear structure and reduces DSBs level proxied from the number of RAD51 and DMC1 foci in *asy3* (50). One attractive hypothesis is that chromosome fragmentation defects result from interhomolog invasion imposed by perturbed ASY1 localization, together with deregulation of RAD51/DMC1 dynamics in *figl1 asy3*. However, the double *figl1 asy4* mutant showed less severe repair defects as only 25% of metaphase I cells were aberrant, with an overall mean bivalent number reduced to 2.84. ASY1 and ASY3 are recruited in *asy4* but form abnormal patchy and lumpy patterns on the chromosome axis, plus ASY1 is not depleted from the synapsed regions (51). We speculate that the presence of ASY1 and ASY3 in *figl1 asy4* would enforce interhomolog invasions as well as counteract FIGL1 functions up to a certain extent, resulting in partly unrepaired breaks and defects in chiasma formation. In our model (Fig.7), ASY proteins would act as positive modulators of interhomolog recombination, while FIGL1 would counteract interhomolog invasions. Lack of both FIGL1 and ASY proteins would thus lead to aberrant meiotic DSB repair.

### Arabidopsis FIGL1 promotes meiotic repair to sisters when interhomolog bias is impaired

In plants, RAD51 repairs meiotic DSBs using sister chromatids in *dmc1* (24, 25), suggesting either plant RAD51 is unable to perform interhomolog invasions or RAD51-mediated interhomolog invasions in *dmc1* can be actively counteracted by an unknown factor(s). Our data show that FIGL1 does not modulate RAD51-mediated intersister repair in *dmc1* or haploid meiosis. We observed univalents and no fragmentation in *Arabidopsis figl1 dmc1* and haploid *figl1* mutant (Fig. 2). However, we noticed the presence of rare bivalents and slightly higher fertility in *figl1 dmc1* compared to *dmc1*. This favors the idea that RAD51 could mediate interhomolog invasions in low frequency in *dmc1* and that FIGL1 counteracts these interhomolog invasions in *dmc1*.

DMC1 is crucial for interhomolog bias, and its dysfunction leads to weakened interhomolog bias in *hop2-2* and *asy1* (34, 49, 83). The insufficient amount of functional HOP2/MND1 complex results in the majority of univalents and DMC1-dependent rare bivalents (∼0.5 per cell) at metaphase I in *hop2-2* (34). Similarly, most univalents and a few DMC1-dependent bivalents (∼1.7 per cell) are observed when the stability of DMC1 is compromised in *asy1*. DMC1 dysfunction thus leads to more repair on sisters at the expense of lower interhomolog repair in *hop2-2* and *asy1*. We found that FIGL1 is critical for this shift in meiotic DSB repair in *hop2-2* and *asy1*. How does FIGL1 pivot between interhomolog and intersister repair? Our data show a hyperaccumulation of RAD51 foci in *figl1 hop2-2* and *figl1 asy1* meiocytes, compared to *hop2-2* and *asy1*, but similar to *figl1*. This indicates that RAD51 foci accumulation largely depends on FIGL1 and is likely independent of HOP2 and ASY1. Since FIGL1 is dispensable for intersister repair (in *dmc1*) but counteracts interhomolog invasions (in *asy1* and *dmc1*), our data imply that FIGL1 dismantles a subset of RAD51 filaments, in addition to DMC1 filaments, arising from interhomolog invasions in *hop2-2* and *asy1* to promote repair on sisters. The strong fragmentation observed in *figl1 hop2-2* and *figl1 asy1* but not in *figl1* also suggests that HOP2 and ASY1 promote DSB repair in *figl1*. We speculate that an interplay between HOP2, ASY1, and FIGL1 is required for wild-type levels of interhomolog recombination in Arabidopsis. Unlike the antagonism between BRCA2 or SDS and FIGL1 at the nucleofilament formation step, we propose that ASY1 could counteract FIGL1 activity for a subset of recombinase foci at the post invasion steps.

## Conclusions

In conclusion, the genetic interaction of *figl1* revealed that FIGL1 attenuates interhomolog repair during meiosis and that these genetic interactions are an essential determinant of the meiotic break repair outcome. We have previously shown that BRCA2 and SDS antagonize FIGL1 activity to protect RAD51/DMC1 foci, likely at the step of nucleofilament formation. This study suggests that multiple factors can counteract FIGL1 activity, in addition to BRCA2 and SDS, likely at the invasion step after nucleofilament formation to promote meiotic interhomolog repair.

## Material and Methods

### Plant growth and genetic material

Arabidopsis plants were cultivated in the greenhouse or growth chamber with 16 h/day and 8h/ night photoperiod at 20°C. *A. thaliana* accession Columbia (Col-0) was the wild type reference. The following Arabidopsis lines were used in this study: *figl1* (*figl1-1*) (57), *rad51*(GABI_134A01), *RAD51-GFP* (28), *mnd1* (SALK_110052), *dmc1* (SAIL_170_F08), *rad51b* (SALK_024755), *xrcc2* (SALK_029106), *asy1* (SALK_046272), *asy3* (SALK_143676), *asy4* (SK22114, CS1006148) (51), *rad54-2* (SALK_124992) (40). Only *hop2-2* allele (EYU48) (34) was used in the background of Wassilewskija (ws) ecotype. Haploid *figl1* plants were produced by crossing Genome Elimination (GEM) lines (84) with the *figl1^+/-^ heterozygous* plants.

### Fertility analysis

Fertility of plants was examined by counting seeds per silique (fruit) on the scanned image of siliques fixed in 70% ethanol using Zeiss Zen software. At least 10 siliques per plant were sampled and sister wild-type plants in the segregating population cultivated together with mutant were used as controls. The unpaired Kruskal-Wallis test corrected by Dunn’s test for multiple comparisons was used to compare the average number of seeds per silique (using Prism GraphPad 9.3.1 software).

### Cytological Techniques

The surface spreads of meiotic chromosomes from pollen mother cells were prepared as previously described (85). Chromosome were visualized by staining with DAPI (1µg/ml). The immunolocalization of MLH1 and HEI10 was performed on male meiocytes with technique that preserves the 3D structure of the meiocyte nucleus, as described in (86). For cytological detection of RAD51, DMC1 and REC8, male meiotic chromosome spreads from prophase I were prepared as described in Armstrong et al. (48). Spread slides were either immediately used for immuno-cytology or stored at -80 C° before immunostaining. Chromosome axis protein REC8 and synaptonemal complex protein ZYP1 staining were performed to identify prophase I cells. The primary antibodies used were: rabbit or rat α-REC8 (1:250) (87), rat α-RAD51 (1:500) (8), rabbit α-DMC1 (1:200) (26), guinea Pig α-ZYP1 (1:250), rabbit α-MLH1(1:1000) (88), chicken α-HEI10 (1:10,000) (71) and rat α-ZYP (89). Secondary antibodies used were conjugated with Alexa Fluor 488 (Thermo Fisher Scientific A27034), Alexa Fluor 568 (Thermo Fisher Scientific A11041), and Alexa Fluor 647 (Thermo Fisher Scientific A21247).

For the chromosome spreads and the 3D immunolocalization, images were acquired using the ZEISS Axio Imager 2 microscope with a 100× oil immersion objective driven by ZEN 2 Software (Carl Zeiss Microscopy). Counting of MLH1-HEI10 co-foci was performed using the Imaris software and analysis of chromosome spreads on ZEN software. Counting of RAD51 and DMC1 foci were computed by Fiji as described previously (60). The unpaired Kruskal-Wallis test corrected by Dunn’s test for multiple comparisons was used for the comparison of (i) the average number of MLH1-HEI10 co-foci per cell and (ii) the average number of bivalents per cell (iii) mean number of RAD51 and DMC1 foci per cell, using Prism GraphPad 9.3.1 software.

### AlphaFold2 predictions

AlphaFold2 predictions for protein complex between XRCC2 and FRBD-FIGL1 were computed through the ColabFold notebook (ColabFold v.1.5.5 and AlphaFold2 v.2.2) using a ColabPro+ plan (https://colab.research.google.com/github/sokrypton/ColabFold/blob/main/AlphaFold2.ipynb). The plDDT, PAE and ipTM scores and graphs were provided directly by this notebook. No template information was used for structure prediction. The predicted structures were not relaxed using amber. mmseqs_uniref_env was used for the unpaired MSA, and sequences from the same species were paired. For the advanced settings, the automatic modes were applied, only one seed was used and dropouts were not enabled. The UCSF ChimeraX (version 1.7.1) was used for producing images in figure.

### Yeast two-hybrid assays

The DNA sequences corresponding to Arabidopsis XRCC2 and FIGL1 flanked by attB1 and attB2 recombinant tails were cloned into pDONOR221 and pDONOR207, respectively, using the Gateway technology (Thermo Fisher). The fidelity of the coding sequence of all clones was verified by sequencing. The entry vectors were used to generate the appropriate pGAD and pGBK yeast two-hybrid expression vectors. Yeast two hybrid assays were carried out using Gal4 based system (Clontech) by introducing plasmids harboring gene of interest in yeast strains AH109 and Y187. After mating of haploid yeasts in YPD plates, diploid cells expressing Gal4BD and Gal4AD fusion proteins were selected in SD/LW, a drop-out medium without leucine and tryptophan. Protein interactions were assayed by growing diploids cells in serial dilutions for 5 days at 30°C on selective media lacking leucine, tryptophan, histidine, and adenine (SD/LWH and SD/LWHA). Two proteins were deemed to interact when the spots grew on SD-LWH and/or SD-LWHA and no self-activation could be observed.

## Acknowledgments

We are grateful to Christine Mezard and Mathilde Grelon for fruitful discussions and critical reading of the manuscript. We also thank Christine Mezard for the technical help in generating Alphafold2 models. We thank Oliver Da Ines for XRCC2 construct and Rabbit anti-DMC1 antibodies.

## Authors contributions

C.E., S.P., J.G., A.H., A.C., and C.G. produced the data. C.E., C.G., R.M. and R.K. analyzed the data. R.K. and R.M. conceived and designed the experiments. C.E. and R.K. wrote the manuscript with the input of all authors.

## Supplemental Figures

**Figure S1.**
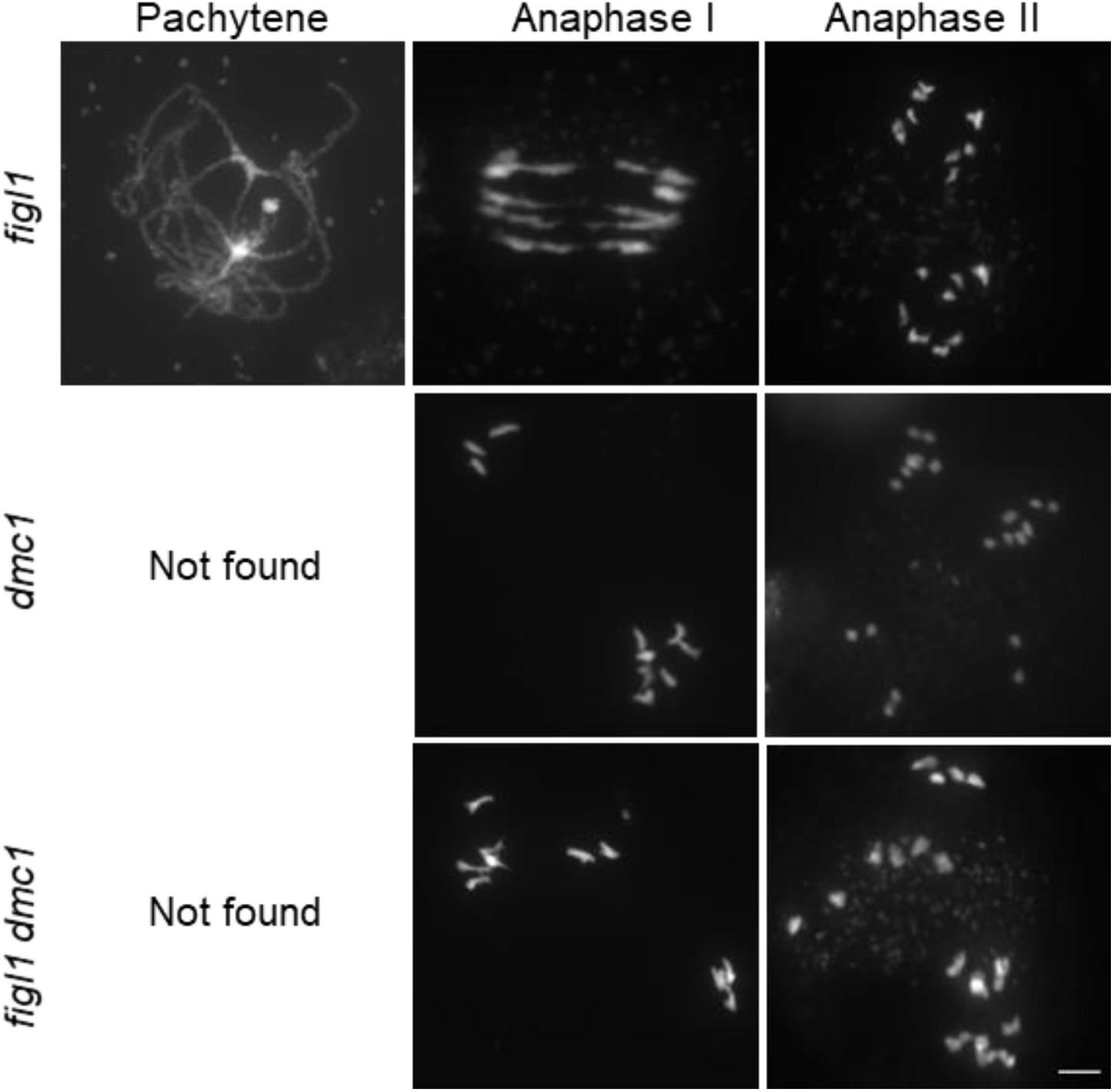
Functional interaction of *figl1* and *dmc1*. Representative Pachytene, Anaphase I, and Anaphase II images of DAPI-stained chromosome spread of male meiocytes are shown in *figl1*, *dmc1,* and *figl1 dmc1.* scale bars: 5µm.

**Figure S2.**
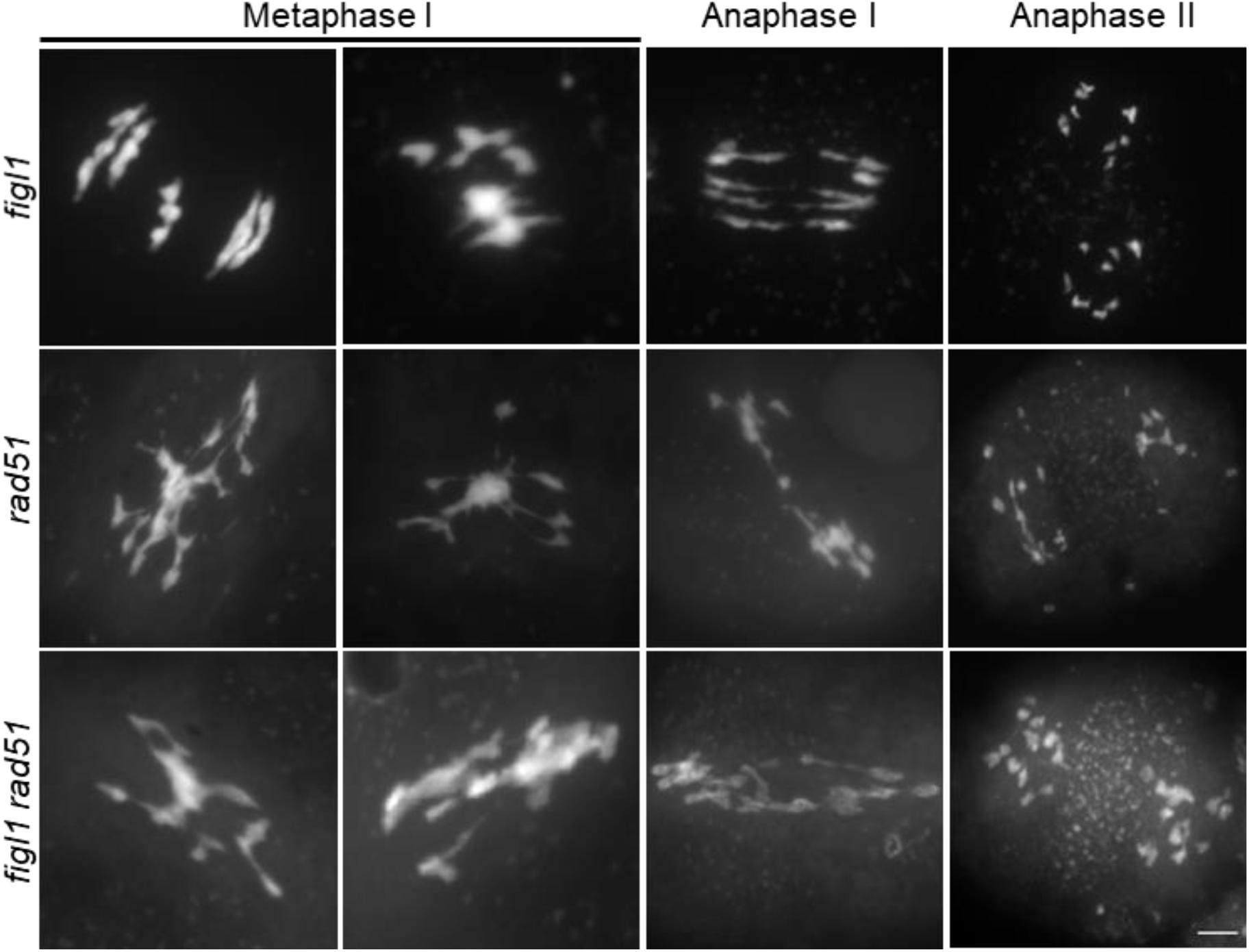
Functional interaction of *figl1* and *rad51*. (A) Representative metaphase I and anaphase II images of DAPI-stained chromosome spread of male meiocytes are shown in *figl1, rad51*, and *figl1 rad51*; scale bars: 5µm

**Fig. S3.**
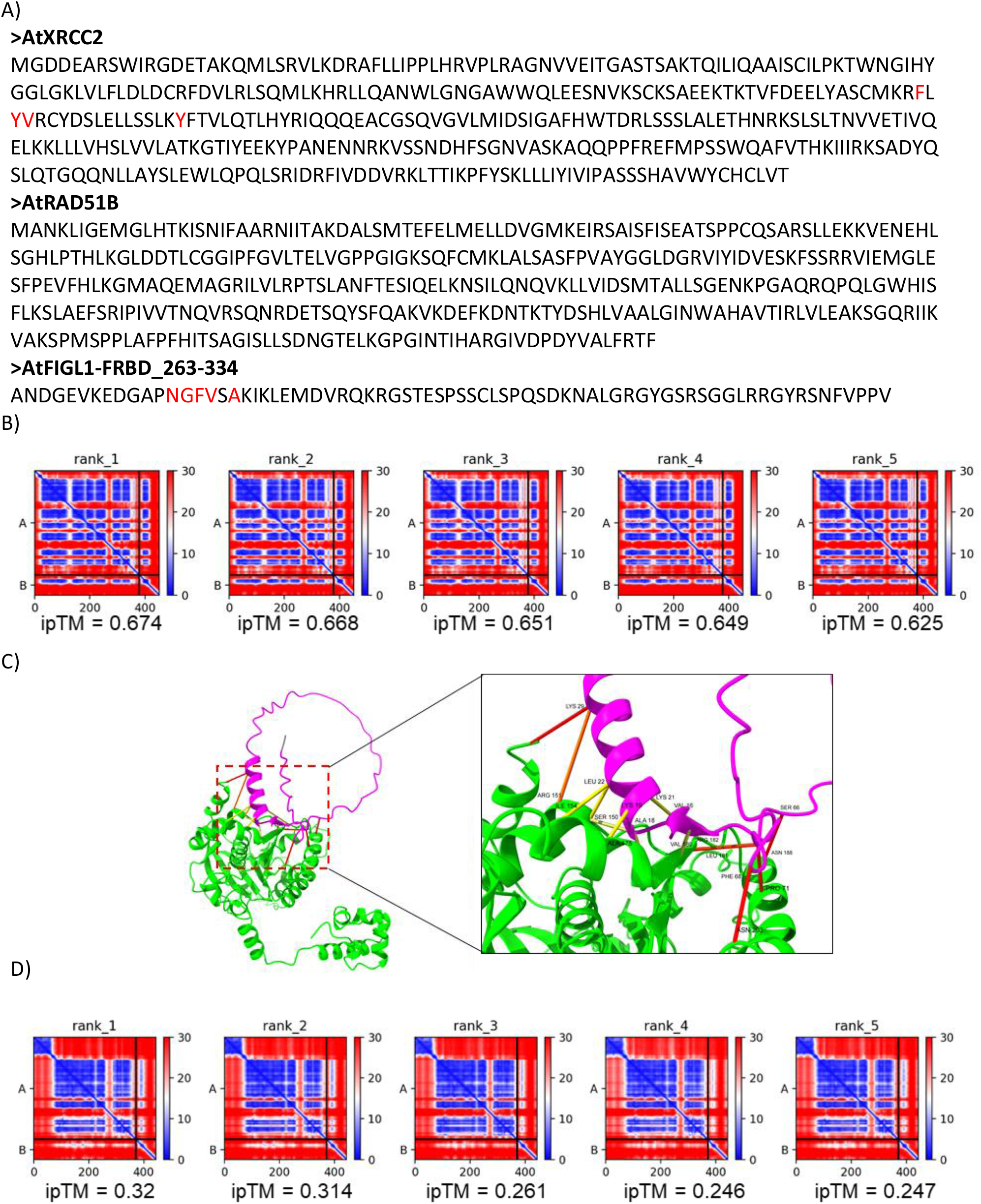
Interaction models of XRCC2, RAD51B and FRBD-FIGL1. A) Sequences of Arabidopsis XRCC2, RAD51B and FIGL-FRBD domain used for 3D structure model prediction by Alphafold2. Residues predicted to be implicated in the interaction are marked in Red. B) Predicted aligned error (PAE) values of five models are shown. Low PAE values in blue indicate strong confidence in the distances between two amino acids, while Red denotes low confidence with high PAE values. A and B denote XRCC2 and FRBD, respectively. C) 3D structure model of interaction between Arabidopsis RAD51B (in green) and FIGL-FRBD domain (in magenta) by Alphafold2. An enlarged 3D view of the binding interface between RAD51B and FIGL1-FRBD domain. Yellow, orange and red lines indicate the low confidence predicted aligned error (PAE) scores for residues shown by Alphafold2. B) Predicted aligned error (PAE) values of five models of RAD51B and FIGL1-FRBD are shown. Medium range PAE values in white indicate not a high confidence in the distances between two amino acids. A and B denote RAD51B and FRBD, respectively.

**Fig. S4.**
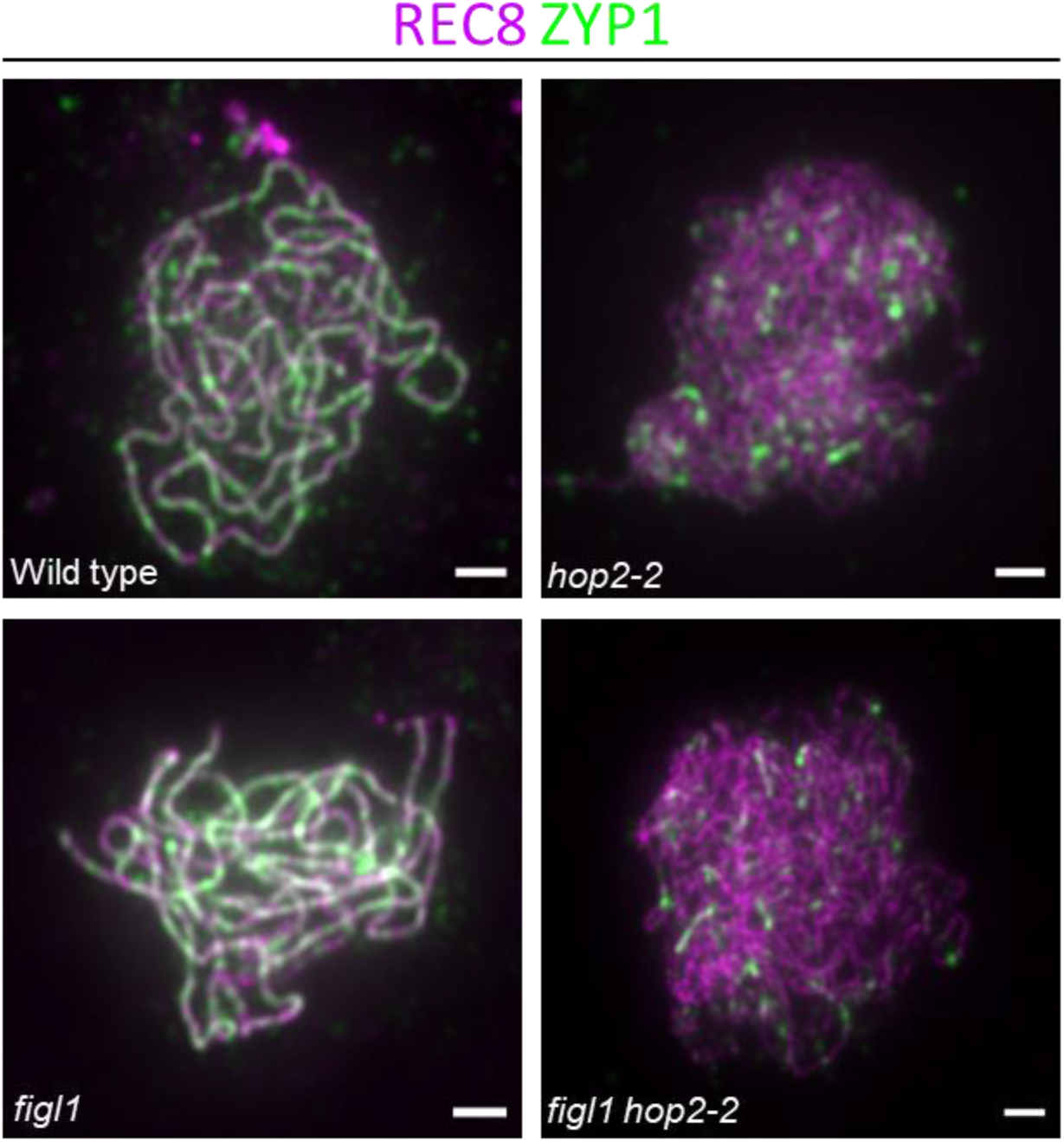
Analysis of synaptonemal complex assembly. Representative images showing dual immunolocalization of REC8 (magenta) and ZYP1 (green) on male meiocytes from wild type, *figl1, hop2-2*, *figl1 hop2-2* in the Col-0/Ws hybrid background. Scale bars: 5µm.

## Bibliography

1. Hunter, N. (2015) Meiotic Recombination: The Essence of Heredity. Cold Spring Harb Perspect Biol, 7, a016618.

2. Wang, Y. and Copenhaver, G.P. (2018) Meiotic Recombination: Mixing It Up in Plants. Annu Rev Plant Biol, 69, 577–609.

3. Schwacha, A. and Kleckner, N. (1994) Identification of joint molecules that form frequently between homologs but rarely between sister chromatids during yeast meiosis. Cell, 76, 51–63.

4. Schwacha, A. and Kleckner, N. (1997) Interhomolog bias during meiotic recombination: meiotic functions promote a highly differentiated interhomolog-only pathway. Cell, 90, 1123–1135.

5. Brown, M.S. and Bishop, D.K. (2014) DNA strand exchange and RecA homologs in meiosis. Cold Spring Harb Perspect Biol, 7, a016659.

6. Humphryes, N. and Hochwagen, A. (2014) A non-sister act: recombination template choice during meiosis. Exp Cell Res, 329, 53–60.

7. Sheridan, S.D., Yu, X., Roth, R., Heuser, J.E., Sehorn, M.G., Sung, P., Egelman, E.H. and Bishop, D.K. (2008) A comparative analysis of Dmc1 and Rad51 nucleoprotein filaments. Nucleic Acids Research, 36, 4057–4066.

8. Kurzbauer, M.-T., Uanschou, C., Chen, D. and Schlögelhofer, P. (2012) The Recombinases DMC1 and RAD51 Are Functionally and Spatially Separated during Meiosis in *Arabidopsis*. Plant Cell, 24, 2058–2070.

9. Brown, M.S., Grubb, J., Zhang, A., Rust, M.J. and Bishop, D.K. (2015) Small Rad51 and Dmc1 Complexes Often Co-occupy Both Ends of a Meiotic DNA Double Strand Break. PLoS Genet, 11, e1005653.

10. Slotman, J.A., Paul, M.W., Carofiglio, F., De Gruiter, H.M., Vergroesen, T., Koornneef, L., Van Cappellen, W.A., Houtsmuller, A.B. and Baarends, W.M. (2020) Super-resolution imaging of RAD51 and DMC1 in DNA repair foci reveals dynamic distribution patterns in meiotic prophase. PLoS Genet, 16, e1008595.

11. Bishop, D.K. (1994) RecA homologs Dmc1 and Rad51 interact to form multiple nuclear complexes prior to meiotic chromosome synapsis. Cell, 79, 1081–1092.

12. Hayashi, M., Chin, G.M. and Villeneuve, A.M. (2007) C. elegans germ cells switch between distinct modes of double-strand break repair during meiotic prophase progression. PLoS Genet, 3, e191.

13. Kim, K.P., Weiner, B.M., Zhang, L., Jordan, A., Dekker, J. and Kleckner, N. (2010) Sister cohesion and structural axis components mediate homolog bias of meiotic recombination. Cell, 143, 924– 937.

14. Crismani, W., Portemer, V., Froger, N., Chelysheva, L., Horlow, C., Vrielynck, N. and Mercier, R. (2013) MCM8 Is Required for a Pathway of Meiotic Double-Strand Break Repair Independent of DMC1 in Arabidopsis thaliana. PLoS Genet, 9, e1003165.

15. Enguita-Marruedo, A., Martín-Ruiz, M., García, E., Gil-Fernández, A., Parra, M.T., Viera, A., Rufas, J.S. and Page, J. (2019) Transition from a meiotic to a somatic-like DNA damage response during the pachytene stage in mouse meiosis. PLoS Genet, 15, e1007439.

16. Toraason, E., Horacek, A., Clark, C., Glover, M.L., Adler, V.L., Premkumar, T., Salagean, A., Cole, F. and Libuda, D.E. (2021) Meiotic DNA break repair can utilize homolog-independent chromatid templates in C. elegans. Curr Biol, 31, 1508–1514.e5.

17. Ziesel, A., Weng, Q., Ahuja, J.S., Bhattacharya, A., Dutta, R., Cheng, E., Börner, G.V., Lichten, M. and Hollingsworth, N.M. (2022) Rad51-mediated interhomolog recombination during budding yeast meiosis is promoted by the meiotic recombination checkpoint and the conserved Pif1 helicase. PLoS Genet, 18, e1010407.

18. Tsubouchi, H. and Roeder, G.S. (2006) Budding yeast Hed1 down-regulates the mitotic recombination machinery when meiotic recombination is impaired. Genes Dev, 20, 1766– 1775.

19. Busygina, V., Sehorn, M.G., Shi, I.Y., Tsubouchi, H., Roeder, G.S. and Sung, P. (2008) Hed1 regulates Rad51-mediated recombination via a novel mechanism. Genes Dev, 22, 786–795.

20. Niu, H., Wan, L., Busygina, V., Kwon, Y., Allen, J.A., Li, X., Kunz, R.C., Kubota, K., Wang, B., Sung, P., et al. (2009) Regulation of meiotic recombination via Mek1-mediated Rad54 phosphorylation. Mol Cell, 36, 393–404.

21. Lao, J.P., Cloud, V., Huang, C.-C., Grubb, J., Thacker, D., Lee, C.-Y., Dresser, M.E., Hunter, N. and Bishop, D.K. (2013) Meiotic Crossover Control by Concerted Action of Rad51-Dmc1 in Homolog Template Bias and Robust Homeostatic Regulation. PLoS Genet, 9, e1003978.

22. Callender, T.L., Laureau, R., Wan, L., Chen, X., Sandhu, R., Laljee, S., Zhou, S., Suhandynata, R.T., Prugar, E., Gaines, W.A., et al. (2016) Mek1 Down Regulates Rad51 Activity during Yeast Meiosis by Phosphorylation of Hed1. PLoS Genet, 12, e1006226.

23. Joshi, N., Brown, M.S., Bishop, D.K. and Börner, G.V. (2015) Gradual implementation of the meiotic recombination program via checkpoint pathways controlled by global DSB levels. Mol Cell, 57, 797–811.

24. Couteau, F., Belzile, F., Horlow, C., Grandjean, O., Vezon, D. and Doutriaux, M.P. (1999) Random chromosome segregation without meiotic arrest in both male and female meiocytes of a dmc1 mutant of Arabidopsis. Plant Cell, 11, 1623–1634.

25. Wang, H., Hu, Q., Tang, D., Liu, X., Du, G., Shen, Y., Li, Y. and Cheng, Z. (2016) OsDMC1 Is Not Required for Homologous Pairing in Rice Meiosis. Plant Physiol, 171, 230–241.

26. Da Ines, O., Bazile, J., Gallego, M.E. and White, C.I. (2022) DMC1 attenuates RAD51-mediated recombination in Arabidopsis. PLoS Genet, 18, e1010322.

27. Cloud, V., Chan, Y.-L., Grubb, J., Budke, B. and Bishop, D.K. (2012) Rad51 is an accessory factor for Dmc1-mediated joint molecule formation during meiosis. Science, 337, 1222–1225.

28. Da Ines, O., Degroote, F., Goubely, C., Amiard, S., Gallego, M.E. and White, C.I. (2013) Meiotic recombination in Arabidopsis is catalysed by DMC1, with RAD51 playing a supporting role. PLoS Genet, 9, e1003787.

29. Seeliger, K., Dukowic-Schulze, S., Wurz-Wildersinn, R., Pacher, M. and Puchta, H. (2012) BRCA2 is a mediator of RAD51- and DMC1-facilitated homologous recombination in *Arabidopsis thaliana*. New Phytologist, 193, 364–375.

30. Fu, R., Wang, C., Shen, H., Zhang, J., Higgins, J.D. and Liang, W. (2020) Rice OsBRCA2 Is Required for DNA Double-Strand Break Repair in Meiotic Cells. Front. Plant Sci., 11, 600820.

31. Azumi, Y. (2002) Homolog interaction during meiotic prophase I in Arabidopsis requires the SOLO DANCERS gene encoding a novel cyclin-like protein. The EMBO Journal, 21, 3081–3095.

32. Petukhova, G.V., Pezza, R.J., Vanevski, F., Ploquin, M., Masson, J.-Y. and Camerini-Otero, R.D. (2005) The Hop2 and Mnd1 proteins act in concert with Rad51 and Dmc1 in meiotic recombination. Nat Struct Mol Biol, 12, 449–453.

33. Chan, Y.-L., Brown, M.S., Qin, D., Handa, N. and Bishop, D.K. (2014) The Third Exon of the Budding Yeast Meiotic Recombination Gene HOP2 Is Required for Calcium-dependent and Recombinase Dmc1-specific Stimulation of Homologous Strand Assimilation. Journal of Biological Chemistry, 289, 18076–18086.

34. Uanschou, C., Ronceret, A., Von Harder, M., De Muyt, A., Vezon, D., Pereira, L., Chelysheva, L., Kobayashi, W., Kurumizaka, H., Schlögelhofer, P., et al. (2014) Sufficient Amounts of Functional HOP2/MND1 Complex Promote Interhomolog DNA Repair but Are Dispensable for Intersister DNA Repair during Meiosis in *Arabidopsis*. The Plant Cell, 25, 4924–4940.

35. Kerzendorfer, C., Vignard, J., Pedrosa-Harand, A., Siwiec, T., Akimcheva, S., Jolivet, S., Sablowski, R., Armstrong, S., Schweizer, D., Mercier, R., et al. (2006) The *Arabidopsis thaliana MND1* homologue plays a key role in meiotic homologous pairing, synapsis and recombination. Journal of Cell Science, 119, 2486–2496.

36. Panoli, A.P., Ravi, M., Sebastian, J., Nishal, B., Reddy, T.V., Marimuthu, M.P.A., Subbiah, V., Vijaybhaskar, V. and Siddiqi, I. (2006) AtMND1 is required for homologous pairing during meiosis in Arabidopsis. BMC Mol Biol, 7, 24.

37. Vignard, J., Siwiec, T., Chelysheva, L., Vrielynck, N., Gonord, F., Armstrong, S.J., Schlögelhofer, P. and Mercier, R. (2007) The interplay of RecA-related proteins and the MND1-HOP2 complex during meiosis in Arabidopsis thaliana. PLoS Genet, 3, 1894–1906.

38. Stronghill, P., Pathan, N., Ha, H., Supijono, E. and Hasenkampf, C. (2010) Ahp2 (Hop2) function in Arabidopsis thaliana (Ler) is required for stabilization of close alignment and synaptonemal complex formation except for the two short arms that contain nucleolus organizer regions. Chromosoma, 119, 443–458.

39. Farahani-Tafreshi, Y., Wei, C., Gan, P., Daradur, J., Riggs, C.D. and Hasenkampf, C.A. (2022) The Arabidopsis HOP2 gene has a role in preventing illegitimate connections between nonhomologous chromosome regions. Chromosome Res, 30, 59–75.

40. Hernandez Sanchez-Rebato, M., Bouatta, A.M., Gallego, M.E., White, C.I. and Da Ines, O. (2021) RAD54 is essential for RAD51-mediated repair of meiotic DSB in Arabidopsis. PLoS Genet, 17, e1008919.

41. Bleuyard, J.-Y., Gallego, M.E., Savigny, F. and White, C.I. (2005) Differing requirements for the Arabidopsis Rad51 paralogs in meiosis and DNA repair. Plant J, 41, 533–545.

42. Osakabe, K., Yoshioka, T., Ichikawa, H. and Toki, S. (2002) Molecular cloning and characterization of RAD51-like genes from Arabidopsis thaliana. Plant Mol Biol, 50, 71–81.

43. Bleuyard, J.-Y. and White, C.I. (2004) The Arabidopsis homologue of Xrcc3 plays an essential role in meiosis. EMBO J, 23, 439–449.

44. Abe, K., Osakabe, K., Nakayama, S., Endo, M., Tagiri, A., Todoriki, S., Ichikawa, H. and Toki, S. (2005) Arabidopsis RAD51C gene is important for homologous recombination in meiosis and mitosis. Plant Physiol, 139, 896–908.

45. Da Ines, O., Degroote, F., Amiard, S., Goubely, C., Gallego, M.E. and White, C.I. (2013) Effects of XRCC2 and RAD51B mutations on somatic and meiotic recombination in Arabidopsis thaliana. Plant J, 74, 959–970.

46. Morgan, C., Nayak, A., Hosoya, N., Smith, G.R. and Lambing, C. (2023) Meiotic chromosome organization and its role in recombination and cancer. In Current Topics in Developmental Biology. Elsevier, Vol. 151, pp. 91–126.

47. Caryl, A.P., Armstrong, S.J., Jones, G.H. and Franklin, F.C. (2000) A homologue of the yeast HOP1 gene is inactivated in the Arabidopsis meiotic mutant asy1. Chromosoma, 109, 62–71.

48. Armstrong, S.J., Caryl, A.P., Jones, G.H. and Franklin, F.C.H. (2002) Asy1, a protein required for meiotic chromosome synapsis, localizes to axis-associated chromatin in Arabidopsis and Brassica. J Cell Sci, 115, 3645–3655.

49. Sanchez-Moran, E., Santos, J.-L., Jones, G.H. and Franklin, F.C.H. (2007) ASY1 mediates AtDMC1- dependent interhomolog recombination during meiosis in Arabidopsis. Genes Dev, 21, 2220– 2233.

50. Ferdous, M., Higgins, J.D., Osman, K., Lambing, C., Roitinger, E., Mechtler, K., Armstrong, S.J., Perry, R., Pradillo, M., Cuñado, N., et al. (2012) Inter-homolog crossing-over and synapsis in Arabidopsis meiosis are dependent on the chromosome axis protein AtASY3. PLoS Genet, 8, e1002507.

51. Chambon, A., West, A., Vezon, D., Horlow, C., De Muyt, A., Chelysheva, L., Ronceret, A., Darbyshire, A., Osman, K., Heckmann, S., et al. (2018) Identification of ASYNAPTIC4, a Component of the Meiotic Chromosome Axis. Plant Physiol, 178, 233–246.

52. Mercier, R., Mézard, C., Jenczewski, E., Macaisne, N. and Grelon, M. (2015) The molecular biology of meiosis in plants. Annu Rev Plant Biol, 66, 297–327.

53. Wang, S., Zickler, D., Kleckner, N. and Zhang, L. (2015) Meiotic crossover patterns: obligatory crossover, interference and homeostasis in a single process. Cell Cycle, 14, 305–314.

54. Lloyd, A. (2023) Crossover patterning in plants. Plant Reprod, 36, 55–72.

55. Crismani, W., Girard, C., Froger, N., Pradillo, M., Santos, J.L., Chelysheva, L., Copenhaver, G.P., Horlow, C. and Mercier, R. (2012) FANCM Limits Meiotic Crossovers. Science, 336, 1588–1590.

56. Girard, C., Crismani, W., Froger, N., Mazel, J., Lemhemdi, A., Horlow, C. and Mercier, R. (2014) FANCM-associated proteins MHF1 and MHF2, but not the other Fanconi anemia factors, limit meiotic crossovers. Nucleic Acids Res, 42, 9087–9095.

57. Girard, C., Chelysheva, L., Choinard, S., Froger, N., Macaisne, N., Lehmemdi, A., Mazel, J., Crismani, W. and Mercier, R. (2015) AAA-ATPase FIDGETIN-LIKE 1 and Helicase FANCM Antagonize Meiotic Crossovers by Distinct Mechanisms. PLoS Genet, 11, e1005369.

58. Séguéla-Arnaud, M., Crismani, W., Larchevêque, C., Mazel, J., Froger, N., Choinard, S., Lemhemdi, A., Macaisne, N., Van Leene, J., Gevaert, K., et al. (2015) Multiple mechanisms limit meiotic crossovers: TOP3α and two BLM homologs antagonize crossovers in parallel to FANCM. Proc. Natl. Acad. Sci. U.S.A., 112, 4713–4718.

59. Séguéla-Arnaud, M., Choinard, S., Larchevêque, C., Girard, C., Froger, N., Crismani, W. and Mercier, R. (2017) RMI1 and TOP3α limit meiotic CO formation through their C-terminal domains. Nucleic Acids Res, 45, 1860–1871.

60. Fernandes, J.B., Duhamel, M., Seguéla-Arnaud, M., Froger, N., Girard, C., Choinard, S., Solier, V., De Winne, N., De Jaeger, G., Gevaert, K., et al. (2018) FIGL1 and its novel partner FLIP form a conserved complex that regulates homologous recombination. PLoS Genet, 14, e1007317.

61. Li, X., Zhang, J., Huang, J., Xu, J., Chen, Z., Copenhaver, G.P. and Wang, Y. (2021) Regulation of interference-sensitive crossover distribution ensures crossover assurance in *Arabidopsis*. Proc. Natl. Acad. Sci. U.S.A., 118, e2107543118.

62. Kumar, R., Duhamel, M., Coutant, E., Ben-Nahia, E. and Mercier, R. (2019) Antagonism between BRCA2 and FIGL1 regulates homologous recombination. Nucleic Acids Res, 47, 5170–5180.

63. Matsuzaki, K., Kondo, S., Ishikawa, T. and Shinohara, A. (2019) Human RAD51 paralogue SWSAP1 fosters RAD51 filament by regulating the anti-recombinase FIGNL1 AAA+ ATPase. Nat Commun, 10, 1407.

64. Zhang, P., Zhang, Y., Sun, L., Sinumporn, S., Yang, Z., Sun, B., Xuan, D., Li, Z., Yu, P., Wu, W., et al. (2017) The Rice AAA-ATPase OsFIGNL1 Is Essential for Male Meiosis. Front. Plant Sci., 8, 1639.

65. Zhang, Q., Fan, J., Xu, W., Cao, H., Qiu, C., Xiong, Y., Zhao, H., Wang, Y., Huang, J. and Yu, C. (2023) The FLIP-FIGNL1 complex regulates the dissociation of RAD51/DMC1 in homologous recombination and replication fork restart. Nucleic Acids Research, 10.1093/nar/gkad596.

66. Yang, S., Zhang, C., Cao, Y., Du, G., Tang, D., Li, Y., Shen, Y., Yu, H. and Cheng, Z. (2022) FIGNL1 Inhibits Non-homologous Chromosome Association and Crossover Formation. Front. Plant Sci., 13, 945893.

67. Ito, M., Furukohri, A., Matsuzaki, K., Fujita, Y., Toyoda, A. and Shinohara, A. (2023) FIGNL1 AAA+ ATPase remodels RAD51 and DMC1 filaments in pre-meiotic DNA replication and meiotic recombination. Nat Commun, 14, 6857.

68. Yuan, J. and Chen, J. (2013) FIGNL1-containing protein complex is required for efficient homologous recombination repair. Proc. Natl. Acad. Sci. U.S.A., 110, 10640–10645.

69. Fernandes, J.B., Séguéla-Arnaud, M., Larchevêque, C., Lloyd, A.H. and Mercier, R. (2018) Unleashing meiotic crossovers in hybrid plants. Proc. Natl. Acad. Sci. U.S.A., 115, 2431–2436.

70. Cifuentes, M., Rivard, M., Pereira, L., Chelysheva, L. and Mercier, R. (2013) Haploid Meiosis in Arabidopsis: Double-Strand Breaks Are Formed and Repaired but Without Synapsis and Crossovers. PLoS ONE, 8, e72431.

71. Chelysheva, L., Vezon, D., Chambon, A., Gendrot, G., Pereira, L., Lemhemdi, A., Vrielynck, N., Le Guin, S., Novatchkova, M. and Grelon, M. (2012) The Arabidopsis HEI10 Is a New ZMM Protein Related to Zip3. PLoS Genet, 8, e1002799.

72. Li, W., Chen, C., Markmann-Mulisch, U., Timofejeva, L., Schmelzer, E., Ma, H. and Reiss, B. (2004) The Arabidopsis AtRAD51 gene is dispensable for vegetative development but required for meiosis. Proc Natl Acad Sci U S A, 101, 10596–10601.

73. Wang, Y., Xiao, R., Wang, H., Cheng, Z., Li, W., Zhu, G., Wang, Y. and Ma, H. (2014) The Arabidopsis RAD51 paralogs RAD51B, RAD51D and XRCC2 play partially redundant roles in somatic DNA repair and gene regulation. New Phytol, 201, 292–304.

74. Mu, N., Li, Y., Li, S., Shi, W., Shen, Y., Yang, H., Zhang, F., Tang, D., Du, G., You, A., et al. (2023) MUS81 is required for atypical recombination intermediate resolution but not crossover designation in rice. New Phytologist, 237, 2422–2434.

75. Charlot, F., Chelysheva, L., Kamisugi, Y., Vrielynck, N., Guyon, A., Epert, A., Le Guin, S., Schaefer, D.G., Cuming, A.C., Grelon, M., et al. (2014) RAD51B plays an essential role during somatic and meiotic recombination in Physcomitrella. Nucleic Acids Res, 42, 11965–11978.

76. Zhang, F., Shen, Y., Miao, C., Cao, Y., Shi, W., Du, G., Tang, D., Li, Y., Luo, Q. and Cheng, Z. (2020) OsRAD51D promotes homologous pairing and recombination by preventing nonhomologous interactions in rice meiosis. New Phytol, 227, 824–839.

77. Zhang, F., Shi, W., Zhou, Y., Ma, L., Miao, Y., Mu, N., Ren, H. and Cheng, Z. (2023) RAD51C-RAD51D interplays with MSH5 and regulates crossover maturation in rice meiosis. New Phytol, 239, 1790–1803.

78. Greenhough, L.A., Liang, C.-C., Belan, O., Kunzelmann, S., Maslen, S., Rodrigo-Brenni, M.C., Anand, R., Skehel, M., Boulton, S.J. and West, S.C. (2023) Structure and function of the RAD51B–RAD51C– RAD51D–XRCC2 tumour suppressor. Nature, 619, 650–657.

79. Rawal, Y., Jia, L., Meir, A., Zhou, S., Kaur, H., Ruben, E.A., Kwon, Y., Bernstein, K.A., Jasin, M., Taylor, A.B., et al. (2023) Structural insights into BCDX2 complex function in homologous recombination. Nature, 619, 640–649.

80. Liu, J., Renault, L., Veaute, X., Fabre, F., Stahlberg, H. and Heyer, W.-D. (2011) Rad51 paralogues Rad55-Rad57 balance the antirecombinase Srs2 in Rad51 filament formation. Nature, 479, 245–248.

81. Thacker, J. (2005) The RAD51 gene family, genetic instability and cancer. Cancer Letters, 219, 125– 135.

82. Braybrooke, J.P., Li, J.-L., Wu, L., Caple, F., Benson, F.E. and Hickson, I.D. (2003) Functional Interaction between the Bloom’s Syndrome Helicase and the RAD51 Paralog, RAD51L3 (RAD51D). Journal of Biological Chemistry, 278, 48357–48366.

83. Lambing, C., Kuo, P.C., Tock, A.J., Topp, S.D. and Henderson, I.R. (2020) ASY1 acts as a dosage-dependent antagonist of telomere-led recombination and mediates crossover interference in Arabidopsis. Proc Natl Acad Sci U S A, 117, 13647–13658.

84. Ravi, M. and Chan, S.W.L. (2010) Haploid plants produced by centromere-mediated genome elimination. Nature, 464, 615–618.

85. Ross, K.J., Fransz, P. and Jones, G.H. (1996) A light microscopic atlas of meiosis in Arabidopsis thaliana. Chromosome Res, 4, 507–516.

86. Hurel, A., Phillips, D., Vrielynck, N., Mézard, C., Grelon, M. and Christophorou, N. (2018) A cytological approach to studying meiotic recombination and chromosome dynamics in Arabidopsis thaliana male meiocytes in three dimensions. Plant J, 95, 385–396.

87. Cromer, L., Jolivet, S., Horlow, C., Chelysheva, L., Heyman, J., De Jaeger, G., Koncz, C., De Veylder, L. and Mercier, R. (2013) Centromeric cohesion is protected twice at meiosis, by SHUGOSHINs at anaphase I and by PATRONUS at interkinesis. Curr Biol, 23, 2090–2099.

88. Chelysheva, L., Grandont, L., Vrielynck, N., le Guin, S., Mercier, R. and Grelon, M. (2010) An easy protocol for studying chromatin and recombination protein dynamics during Arabidopsis thaliana meiosis: immunodetection of cohesins, histones and MLH1. Cytogenet Genome Res, 129, 143–153.

89. Higgins, J.D., Sanchez-Moran, E., Armstrong, S.J., Jones, G.H. and Franklin, F.C.H. (2005) The Arabidopsis synaptonemal complex protein ZYP1 is required for chromosome synapsis and normal fidelity of crossing over. Genes Dev, 19, 2488–2500.

